# The Neural Blueprint of Stress Susceptibility: Brain-wide neuronal activity associated with the consequences of stress

**DOI:** 10.1101/2024.09.27.615412

**Authors:** Bart C.J. Dirven, Moritz Negwer, Andriana Botan, Riv Maas, Lennart van Melis, Sanne Merjenburgh, Rebecca van Rijn, Joanes Grandjean, Judith R. Homberg, Tamas Kozicz, Marloes J.A.G. Henckens

**Author notes:** Equal contributions (Andriana Botan, Riv Maas, Lennart van Melis, Sanne Merjenburgh, Rebecca van Rijn). Equal contributions (Joanes Grandjean, Judith R. Homberg, Tamas Kozicz).

## Abstract

Understanding the neurobiological mechanisms of stress susceptibility is key to advancing our insight into stress-related psychopathology like post-traumatic stress disorder (PTSD). Preclinical animal models however typically lack translationally relevant brain readouts. Here, we used a mouse model for severe stress, segregating mice into stress-susceptible and resilient groups. We analyzed and contrasted their whole-brain neuronal activity pre-, peri- and post-stress exposure and compared functional connectivity of the salience (SN), default mode (DMN) and executive control network (lateral cortical network (LCN) in rodents). We found that stress-susceptible mice exhibited pre-existing hyperactivity in the lateral orbital area, a potential risk factor for stress vulnerability, and heightened retrosplenial cortex activity peri- and post-stress, potentially contributing to maladaptive fear memory. Upon stress exposure, susceptible mice showed strong recruitment of visual and memory-related areas, whereas resilient mice displayed marked reductions in retrosplenial activity and increased activation in the agranular insula and ventral striatum. Observations of enhanced intra-SN and SN-DMN connectivity in susceptible mice mirrored those observed in individuals with PTSD. Moreover, susceptible mice displayed aberrant DMN-LCN network pre- and peri-stress. These findings highlight the importance of dynamic network interactions in stress susceptibility and suggest novel brain region targets for (early) intervention.

## Introduction

Every individual has to cope with stressful situations during their lifetime. In susceptible people this may lead to the development of stress-related disorders^1,2^, but most people are able to successfully adapt in the face of severe stress and are resilient to its long-term deleterious effects^3,4^. Improved understanding of the biological basis of interindividual differences in stress responses^5^ is key to enhance insight into stress-related pathophysiology and to generate new leads for their improved prevention, detection, and treatment. Preclinical animal models are instrumental for this, as they allow for tight experimental control, as well as invasive measurements and manipulations at superior spatial resolution and temporal precision. These models increasingly acknowledge the relevance of incorporating interindividual differences in stress resilience/susceptibility to enhance the translational value of the results derived^4,6,7^. Contrasting resilient vs susceptible animals enables one to dissociate adaptive stress responses contributing to full recovery from those that are maladaptive and causative to long-term behavioral symptoms accumulating in stress-related mental disease. Importantly, animal models allow for the study of responses to adversity of an intensity that actually causes such long-term behavioral effects, unlike research using human subjects, which is – for obvious ethical reasons – restricted to studying the responses to mild stressors in rather artificial laboratory settings. Yet, the vast majority of the animal work has focused on isolated brain regions or neuronal circuits of interest, lagging behind on advancing clinical insights implicating disruptions in large-scale functional networks in psychopathology^8^.

Here, we used transgenic (ArcTRAP) mice^9^ that allow for the *in vivo* identification of active neurons across the whole brain at specific time points. Mice were tested in a preclinical model for stress susceptibility^10^, dissociating stress susceptible from resilient mice, and their brain-wide neuronal activity pre-, peri-, and post-stress exposure was compared. This temporal mapping enabled the critical dissociation of deviations reflecting pre-existing risk factors, aberrant acute stress responses, or inadequate stress recovery, carrying unique implications for prevention, early intervention, and treatment of stress-related psychopathology, respectively. Neuronal activity throughout intact brain hemispheres was visualized by immunolabeling-enabled three-dimensional imaging of solvent-cleared organs (iDISCO+)^11^ combined with light-sheet microscopy and activity patterns were compared across groups. Moreover, to assess network functional connectivity, we explored cross-animal activity correlations between brain regions included in the key networks implicated in human psychopathology^8^; the default mode (DMN), salience (SN), and lateral cortical network (LCN; the rodent orthologue of the human executive control network) in resilient and susceptible groups^12^. The findings provide unique insight in the brain-wide spatial and temporal profiles through which aberrant neuronal activity and network connectivity encode stress resilience or susceptibility, paving pave the way towards future mechanistic investigations of these deviations.

## Results

### Interindividual differences in stress susceptibility

The goal of this study was to elucidate the brain-wide neuronal activity patterns and network function associated with stress susceptibility in mice. To distinguish pre-existing differences between stress susceptible and resilient mice conferring mere risk for later symptoms, from acquired deviations, more closely related to the actual behavioral symptoms (or tentatively human pathology), we investigated brain activity at three different timepoints; pre-, peri- or post-stress. Specifically, three cohorts of 44-48 male ArcCreERT2xtdTomato (i.e., ArcTRAP) mice were exposed to a previously established protocol for stress induction by repeated footshock exposure^10,13^, and neuronal activity was assessed in these cohorts at different timepoints by the timed injection of tamoxifen either under home cage conditions pre-stress (cohort 1: n = 48) and post-stress (cohort 3: n = 48) – to mimic assessments of resting-state activity in patients – or peri-stress (cohort 2: n = 44) to assess acute stress responses (Fig. 1a). Tamoxifen injection in these mice induced the indelible labelling of neurons expressing the immediate early gene *Arc*, indicative of neuronal activity, by the fluorescent marker tdTomato. Starting from 1 week post-stress onwards, all mice were behaviorally phenotyped to assess the long-term consequences of stress exposure, focusing on symptoms resembling those observed in PTSD patients, i.e., impaired risk assessment, heightened anxiety, hypervigilance, disrupted pre-pulse-inhibition and sleep disruptions (Fig. 1a). Instead of focusing on single behavioral readouts, we created composite scores (i.e., overall stress symptom scores) by assigning points to the mice displaying the most extreme behavior per readout (see Methods section for further details). Mice displaying multiple of these extreme behaviors were categorized as stress susceptible, whereas mice displaying none were coined resilient^10,13^. We adopted this approach as it more closely resembles the situation in patients, which can be diagnosed with PTSD based on a plethora of (combinations of) symptoms (DSM-V^14^), resulting in high symptom heterogeneity^15^. We similarly observed high variance in behavioral symptoms of mice categorized as susceptible in this mouse model, both within and across experimental cohorts (Fig. 1, see Fig. S1 and Table S1 for the raw data and effect sizes). However, we also saw large differences in stress symptom score compared to resilient groups (Fig. 1b).

**Fig. 1.**
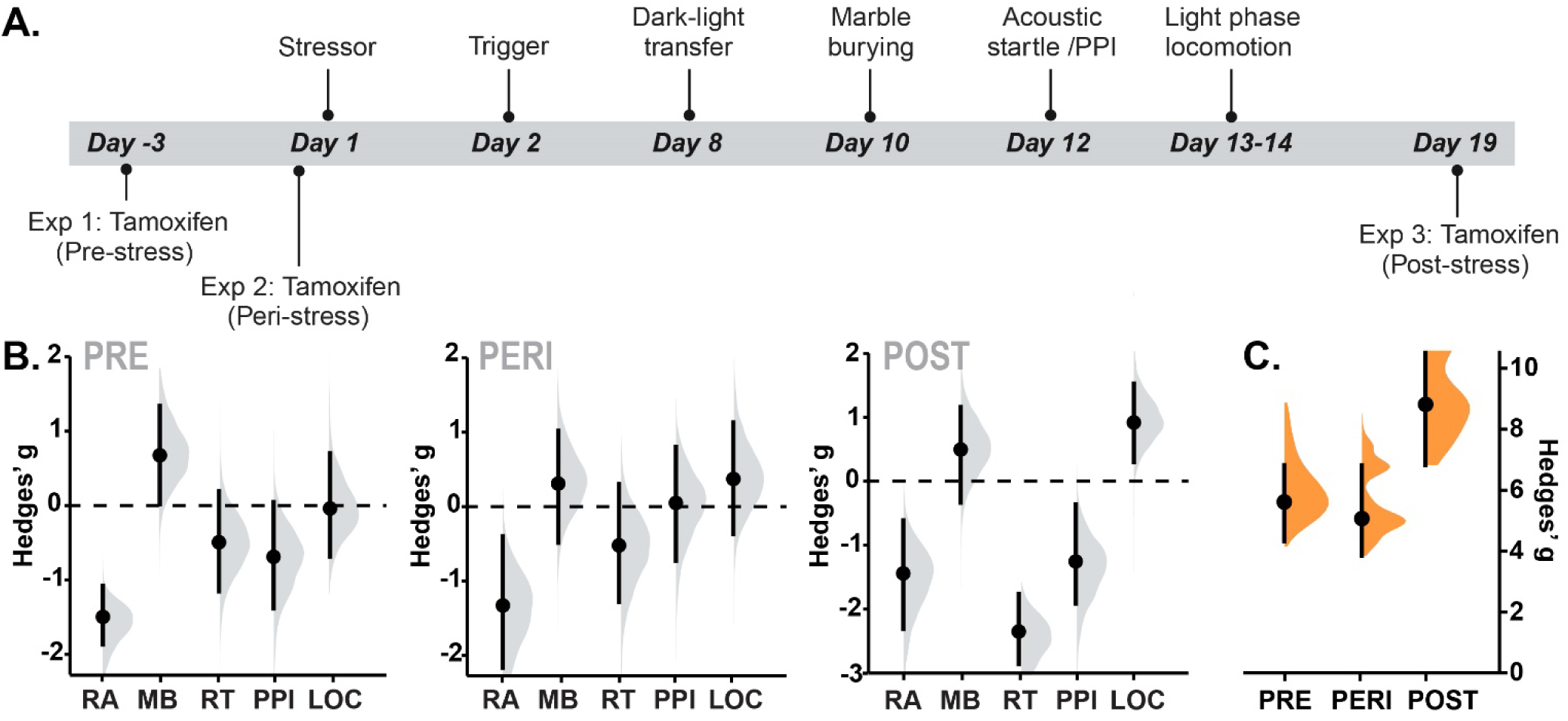
Experimental timeline and behavioral results for the three experiments in which neuronal activity was tagged either pre-stress (experiment 1), peri-stress (experiment 2) or post-stress (experiment 3). Following stress exposure, mice were subjected to several behavioral tasks to assess their risk assessment (RA), marble burying (MB), reaction time (RT) to peak startle response, pre-pulse inhibition (PPI) and locomotor (LOC) activity during the inactive/light phase as symptoms of stress susceptibility (**A**). Mice belonging to the top 20% of each of these behaviors was assigned a score, which were tallied up to generate an overall symptom score; mice with a symptom score of zero were coined resilient, whereas those with a score >= 4 were coined susceptible. Hedges’ g for the difference susceptible > resilient mice for all individual behaviors across experimental cohorts is depicted in Gardner-Altman estimation plots^65^ (**B**). All three behavioral cohorts displayed clear differences in overall symptoms scores between susceptible and resilient mice (**C**). Mean differences (susceptible – resilient) are depicted as a bootstrap sampling distribution; the mean difference is indicated with a dot, whereas the 90% confidence interval is indicated by the ends of the vertical error bars. See Fig. S1 for all raw individual data. Fig. 1a was created with BioRender.com.

### Neuronal activity patterns distinguish susceptible from resilient mice

To investigate brain-wide neuronal activity in these susceptible and resilient mice, they were sacrificed at the end of the paradigm, their brains were extracted, and their brain hemispheres were separated. Left hemispheres were then stained for tdTomato with iDISCO+ and imaged by light-sheet microscopy, allowing for the assessment of activated neurons across the intact hemisphere. The right hemisphere was sliced for dedicated immunohistochemistry studies^16,17^. The 3D light-sheet stacks were mapped on the Allen Brain Atlas and cell counts within 92 anatomically defined regions were analyzed (Table S2). To correct for differences in overall cell detection caused by slight differences in the quality of brain clearing and imaging, regional counts of activated neurons were normalized to the total cell count of the animal’s brain to assess differences in relative distribution of neuronal activity across groups.

Pre-stress (n_resilient_ = 10, n_susceptible_ = 6, Fig. 2a), susceptible mice displayed higher relative cell counts in specifically the lateral orbital area (g_sus>res_ = 1.26, 90% CI = [0.35, 2.13]), ventral striatum (g_sus>res_ = 0.88 [0.02, 1.72]) and medial pallidum (g_sus>res_ = 1.11 [0.22, 1.96]), and lower relative cell counts in the ventral anterior cingulate cortex (g_sus>res_ = -0.86 [-1.70, 0.00]), ventral retrosplenial area (g_sus>res_ = -0.96 [-1.80, - 0.09]), dorsal hippocampal field CA2 (g_sus>res_ = -0.87 [-1.70, 0.00]) and ventral group of the dorsal thalamus (g_sus>res_ = -1.17 [-2.04, -0.27]), compared to resilient animals.

**Fig. 2.**
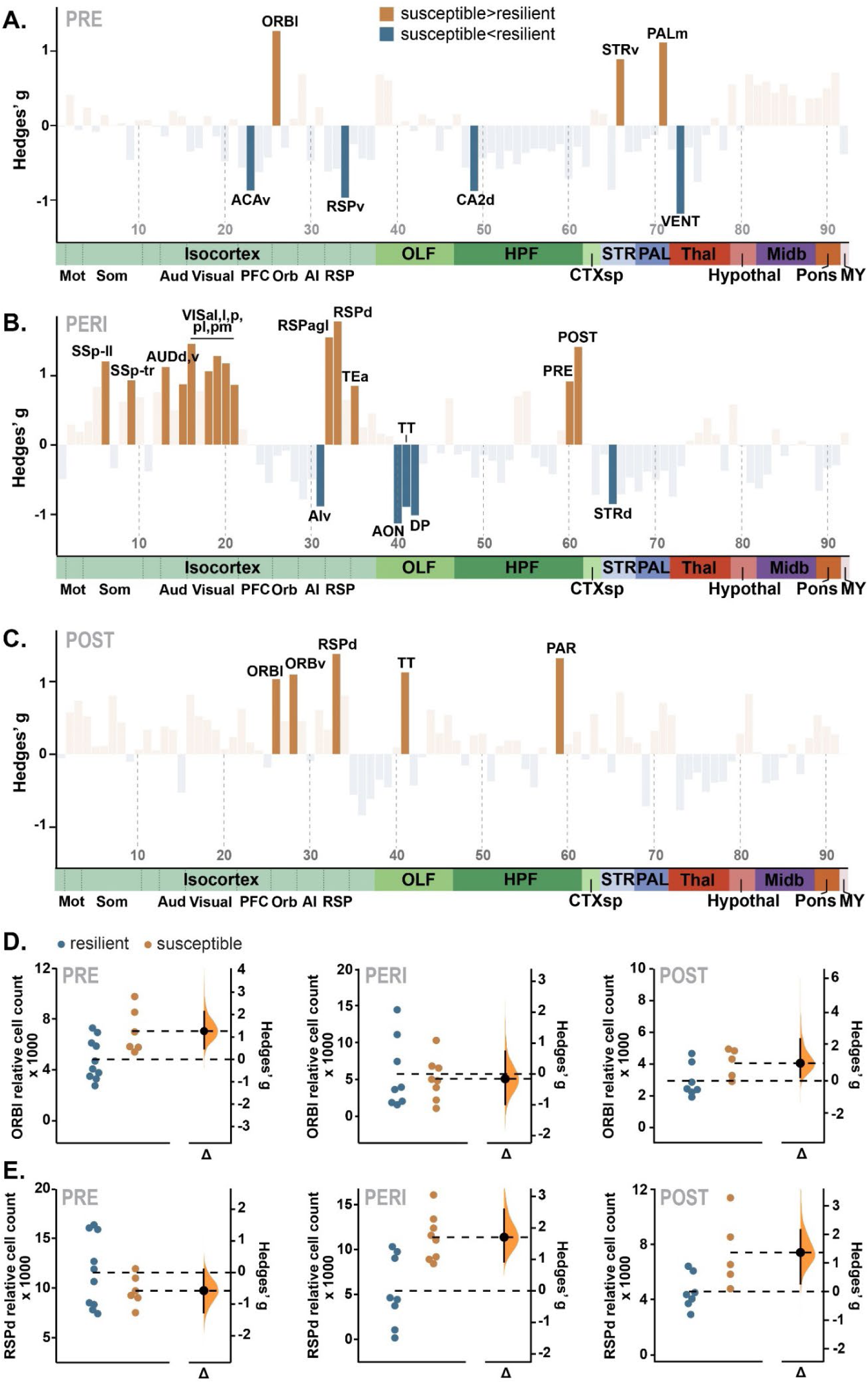
Differences in neuronal activity across the brain between susceptible and resilient mice pre- (**A**), peri- (**B**) and post-stress (**C**). The assessment of differences in neuronal activity across groups at different time points allowed for the identification of pre-existing differences in neuronal activity conferring risk to later stress-related symptoms (e.g., in the lateral orbital area (ORBl) (**D**)), as well as differences acquired over the course of stress exposure (e.g., in the dorsal retrosplenial area (RSPd) (**E**)). A list of the depicted brain regions can be found in Table S2. Opaque bars in panels A-C indicate the differences in normalized cell counts where the 90% confidence interval does not cross or contain 0. Panels D-E show the Hedges’ g for the difference susceptible > resilient mice in Gardner-Altman estimation plots^65^. Both groups are plotted on the left axes; the mean difference is plotted on floating axes on the right as a bootstrap sampling distribution. The mean difference is depicted as a dot; the 90% confidence interval is indicated by the ends of the vertical error bar. ACAv: ventral anterior cingulate area, AIv: ventral agranular insula, AUDd: dorsal auditory area, AUDv: ventral auditory area, AON: anterior olfactory nucleus, CA2d: dorsal hippocampal field CA2, CTXsp: cortical subplate, DP: dorsal peduncular area, HPF: hippocampal formation, Hypothal: hypothalamus, Midb: midbrain, Mot: motor areas, MY: medulla, OLF: olfactory areas, ORBv: ventral orbital area, PALm: medial pallidum, PAR: parasubiculum, PFC: prefrontal cortex, POST: postsubiculum, PRE: presubiculum, RSPagl: lateral agranular retrosplenial area, RSPd: dorsal retrosplenial area, RSPv: ventral retrosplenial area, Som: somatosensory areas, SSp-ll: primary somatosensory area, lower limb, SSp- tr: primary somatosensory area, trunk, STRd: dorsal striatum, STRv: ventral striatum, TEa: temporal association areas, Thal: thalamus, TT: Taenia tecta, VENT: ventral group of the dorsal thalamus, VISal: anterolateral visual area, VISl: lateral visual area, VISp: posterior visual area, VISpl: posterolateral visual area, VISpm: posteromedial visual area.

Peri-stress (n_resilient_ = 9, n_susceptible_ = 8, Fig. 2b), susceptible mice displayed increased relative cell counts compared to resilient mice in primary somatosensory areas (lower limb area: g_sus>res_ = 1.20 [0.32, 2.04], trunk area: g_sus>res_ = 0.93 [0.08, 1.74]), auditory areas (dorsal auditory area: g_sus>res_ = 1.12 [0.25, 1.95], ventral auditory area: g_sus>res_ = 0.87 [0.03, 1.68]), visual areas (anterolateral visual area: g_sus>res_ = 1.45 [0.53, 2.33], lateral visual area: g_sus>res_ = 1.06 [0.20, 1.88], primary visual area: g_sus>res_ = 1.28 [0.39, 2.13], posterolateral visual area: g_sus>res_ = 1.17 [0.30, 2.00], posteromedial visual area: g_sus>res_ = 0.86 [0.03, 1.67]), retrosplenial areas (lateral agranular retrosplenial area: g_sus>res_ = 1.55 [0.61, 2.44], and dorsal retrosplenial area: g_sus>res_ = 1.71 [0.74, 2.62]), subiculum (postsubiculum: g_sus>res_ = 0.91 [0.07, 1.72], presubiculum: g_sus>res_ = 1.41 [0.50, 2.28]) and temporal association areas (g_sus>res_ = 0.85 [0.01, 1.65]). In contrast, the olfactory region (anterior olfactory nucleus (g_sus>res_ = -1.13 [-1.97, -0.26]), taenia tecta (g_sus>res_ = -0.89 [-1.70, -0.05]), dorsal peduncular area (g_sus>res_ = -1.02 [-1.84, -0.16])), ventral agranular insular area (g_sus>res_ = -0.89 [-1.70, -0.05]) and dorsal striatum (g_sus>res_ = -0.85 [-1.66, -0.02]) were characterized by lower relative cell counts in susceptible compared to resilient animals.

Post-stress (n_resilient_ = 7, n_susceptible_ = 5, Fig. 2c), susceptible animals showed higher relative cell counts in orbital areas (lateral orbital area: g_sus>res_ = 1.02 [0.04, 1.97], ventral orbital area: g_sus>res_ = 1.09 [0.09, 2.03]), the taenia tecta (g_sus>res_ = 1.12 [0.12, 2.07]), dorsal retrosplenial area (g_sus>res_ = 1.37 [0.32, 2.36]) and the parasubiculum (g_sus>res_ = 1.31 [0.27, 2.29]) compared to their resilient counterparts.

Interestingly, increased cell counts (indicative of heightened activity) in the lateral orbitofrontal area were observed in both pre- and post-stress cohorts, and therefore identified as potential pre-existing risk factor, or biomarker, for stress susceptibility (Fig. 2d). Higher cell counts in the dorsal retrosplenial area were first observed in susceptible mice during stress exposure and was retained post-stress, potentially reflecting an acquired anomaly related to pathology (Fig. 2e). Noteworthy, voxel-wise comparisons of the group cell density heat maps (Fig. S2), allowing the detection of sub-area clusters, generated similar results as those obtained from the extracted counts.

### Responses to stress in susceptible and resilient mice

Neuronal activity distribution mainly deviated between susceptible vs resilient mice during acute stress exposure. Yet, from this contrast one can impossibly deduce whether the observed differences are caused by either exaggerated, (proposedly) maladaptive responses, or diminished adaptive responses to stress in susceptible mice. To dissociate these two scenarios, we compared neuronal activity during stress exposure to that before stress exposure, to estimate the magnitude of the stress response in susceptible and resilient subgroups. Our analyses firstly revealed clear main effects of stress exposure (Fig. 3a), with increases in relative cell counts in visual areas, agranular insula, piriform, ventral dentate gyrus, entorhinal areas, subiculum and medulla being observed in both groups (see Table S3 for effect sizes and confidence intervals). Conversely, we found relative decreases in cell counts under stress in the primary motor area, primary somatosensory areas, dorsal and ventral thalamic areas and midbrain regions. Yet, of main interest were the stress exposure x group interaction effects observed (Fig. 3b, Table S4). Susceptible mice displayed increases in relative cell counts peri-stress that were not observed in resilient mice in several visual areas (anterolateral (susceptible: g_peri>pre_ = 1.65 [0.44, 2.81], resilient: g_peri>pre_ = -0.25 [-1.14, 0.64]), lateral (susceptible: g_peri>pre_ = 2.59 [1.14, 3.99], resilient: g_peri>pre_ = 0.68 [-0.25, 1.58]) and primary (susceptible: g_peri>pre_ = 2.98 [1.42, 4.50], resilient: g_peri>pre_ = 0.45 [-0.46, 1.34])), temporal association areas (susceptible: g_peri>pre_ = 1.29 [0.15, 2.38], resilient: g_peri>pre_ = 0.39 [-0.51, 1.27]), and post- (susceptible: g_peri>pre_ = 2.10 [0.78, 3.36], resilient: g_peri>pre_ = 0.11 [-0.77, 0.99]) and presubiculum (susceptible: g_peri>pre_ = 2.13 [0.81, 3.41], resilient: g_peri>pre_ = 0.17 [-0.72, 1.05]). Whereas stress-related increases in relative cell counts in the posterolateral visual area (susceptible: g_peri>pre_ = 2.18 [0.84, 3.47], resilient: g_peri>pre_ = 1.33 [0.31, 2.30]) and ventral dentate gyrus (susceptible: g_peri>pre_ = 1.65 [0.44, 2.82], resilient: g_peri>pre_ = 1.09 [0.12, 2.04]) were observed in both groups, these increases were also more pronounced in susceptible mice. Contrarily, stress-related increases in relative cell counts in the dorsal (susceptible: g_peri>pre_ = 0.15 [-0.84, 1.14], resilient: g_peri>pre_ = 1.46 [0.42, 2.46]) and ventral agranular insular area (susceptible: g_peri>pre_ = 0.57 [-0.46, 1.57], resilient: g_peri>pre_ = 1.63 [0.56, 2.66]) and ventral striatum (susceptible: g_peri>pre_ = 0.39 [-0.62, 1.38], resilient: g_peri>pre_ = 1.99 [0.85, 3.09]) were only observed in resilient mice.

**Fig. 3.**
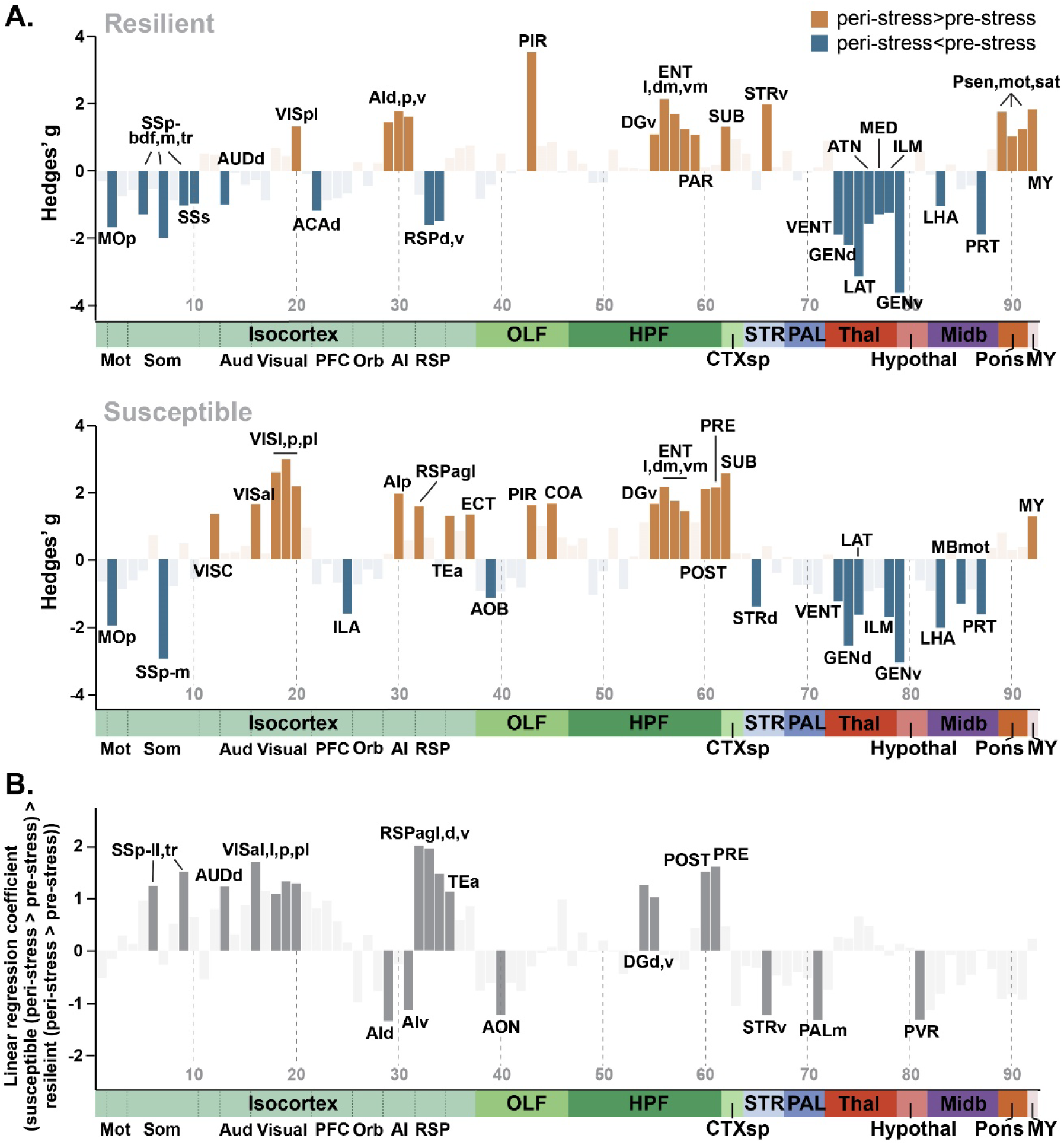
Neuronal activity changes observed under stress in resilient and susceptible mice, and their contrast. Differences in normalized cell counts, indicative of regional neuronal activity, observed in the peri- vs pre- stress cohorts in resilient and susceptible mice, respectively (**A**). Differences in the stress-related changes in neuronal activity between susceptible and resilient mice (**B**). A list of the depicted brain regions can be found in Table S2. Opaque bars indicate the differences in normalized cell counts where the 90% confidence interval does not cross or contain 0. ACAd: dorsal anterior cingulate area, AId: dorsal agranular insula, AIp: posterior agranular insula, AIv: ventral agranular insula, AOB: accessory olfactory bulb, AON: anterior olfactory nucleus, ATN: anterior group of the dorsal thalamus, AUDd: dorsal auditory area, COA: cortical amygdalar area, DGd: dorsal dentate gyrus, DGv: ventral dentate gyrus, ECT: ectorhinal area, ENTdm: medial entorhinal area, dorsal zone, ENTl: lateral entorhinal area, ENTvm: medial entorhinal area, ventral zone, GENd: geniculate group of the dorsal thalamus, GENv: geniculate group of the ventral thalamus, HPF: hippocampal formation, Hypothal: hypothalamus, ILA: infralimbic area, ILM: intralaminar nuclei of the dorsal thalamus, LAT: lateral group of the dorsal thalamus, LHA: lateral hypothalamic area, MBmot: midbrain, motor related, MED: medial group of the dorsal thalamus, Midb: midbrain, MOp: primary motor area, Mot: motor areas, MY: medulla, OLF: olfactory areas, PALm: medial pallidum, PAR: parasubiculum, PFC: prefrontal cortex, PIR: piriform area, Pmot: pons, motor related, POST: postsubiculum, PRE: presubiculum, PRT: pretectal region, Psat: pons, behavioral state related, Psen: pons, sensory related, PVR: periventricular region, RSPagl: lateral agranular retrosplenial area, RSPd: dorsal retrosplenial area, RSPv: ventral retrosplenial area, Som: somatosensory areas, SSp-bdf: primary somatosensory area, barrel field, SSp-ll: primary somatosensory area, lower limb, SSp-m: primary somatosensory area, mouth, SSp-tr: primary somatosensory area, trunk, SSs: supplemental somatosensory area, STRd: dorsal striatum, STRv: ventral striatum, SUB: subiculum, TEa: temporal association areas, Thal: thalamus, VENT: ventral group of the dorsal thalamus, VISal: anterolateral visual area, VISC: visceral area, VISl: lateral visual area, VISp: posterior visual area, VISpl: posterolateral visual area.

Furthermore, reductions in relative cell counts in retrosplenial areas were more pronounced in resilient mice. Resilient mice showed stress-related decreases in relative cell counts in the dorsal (g_peri>pre_ = -1.61 [- 2.63, -0.55]) and ventral (g_peri>pre_ = -1.48 [-2.48, -0.44]) retrosplenial areas, which were absent in susceptible mice (dorsal: g_peri>pre_ = 0.64 [-0.40, 1.65], ventral retrosplenial: g_peri>pre_ = -0.28 [-1.27, 0.72]). Susceptible mice even displayed enhanced activity in the agranular retrosplenial cortex under stress (g_peri>pre_ = 1.58 [0.38, 2.73]), whereas no such effect was seen in resilient mice (g_peri>pre_ = -0.72 [-1.63, 0.21). Additionally, resilient mice displayed reduced relative cell counts in the primary somatosensory trunk area (g_peri>pre_ = - 1.03 [-1.97, -0.06]) and dorsal auditory area (g_peri>pre_ = -1.00 [-1.93, -0.03]), whereas susceptible mice did not (somatosensory trunk: g_peri>pre_ = 0.48 [-0.54, 1.48], dorsal auditory: g_peri>pre_ = 0.17 [-0.82, 1.16]). Finally, the primary somatosensory limb area, anterior olfactory nucleus, dorsal dentate gyrus, medial pallidum, and periventricular region displayed stress exposure x group interaction effects (Table S4). As such, several differences were observed in stress-induced neuronal activation between susceptible and resilient mice, with susceptible mice showing exaggerated (i.e., quantitively different) neuronal activation in some regions, but also a qualitatively different response in others.

### Neuronal activity correlation differences reflect altered neural network connectivity associated with stress susceptibility

Human neuroimaging work has indicated that the brain is organized as a set of large-scale functional networks, each carrying out specialized functions^18^, and that deficits in their recruitment and (dis)engagement might underly psychopathology^19^. Specifically, the triple network model of psychopathology^20^ posits that aberrant function of the salience network (SN), executive control network (ECN), and default mode network (DMN) and their dynamic cross-network interactions encode a wide range of pathologies. Importantly, these networks have also been identified in the rodent brain^21^, with the LCN displaying anti-correlated activity with the DMN^21-23^, resembling the human ECN. Therefore, we next focused our analyses on the differences between susceptible and resilient mice in the activity and connectivity of these networks. Our definition of the DMN, SN and LCN was based on prior viral tracer studies targeting core regions within these networks^24-26^ identifying their structural connectivity (Fig. 4a, Table S5).

**Fig. 4.**
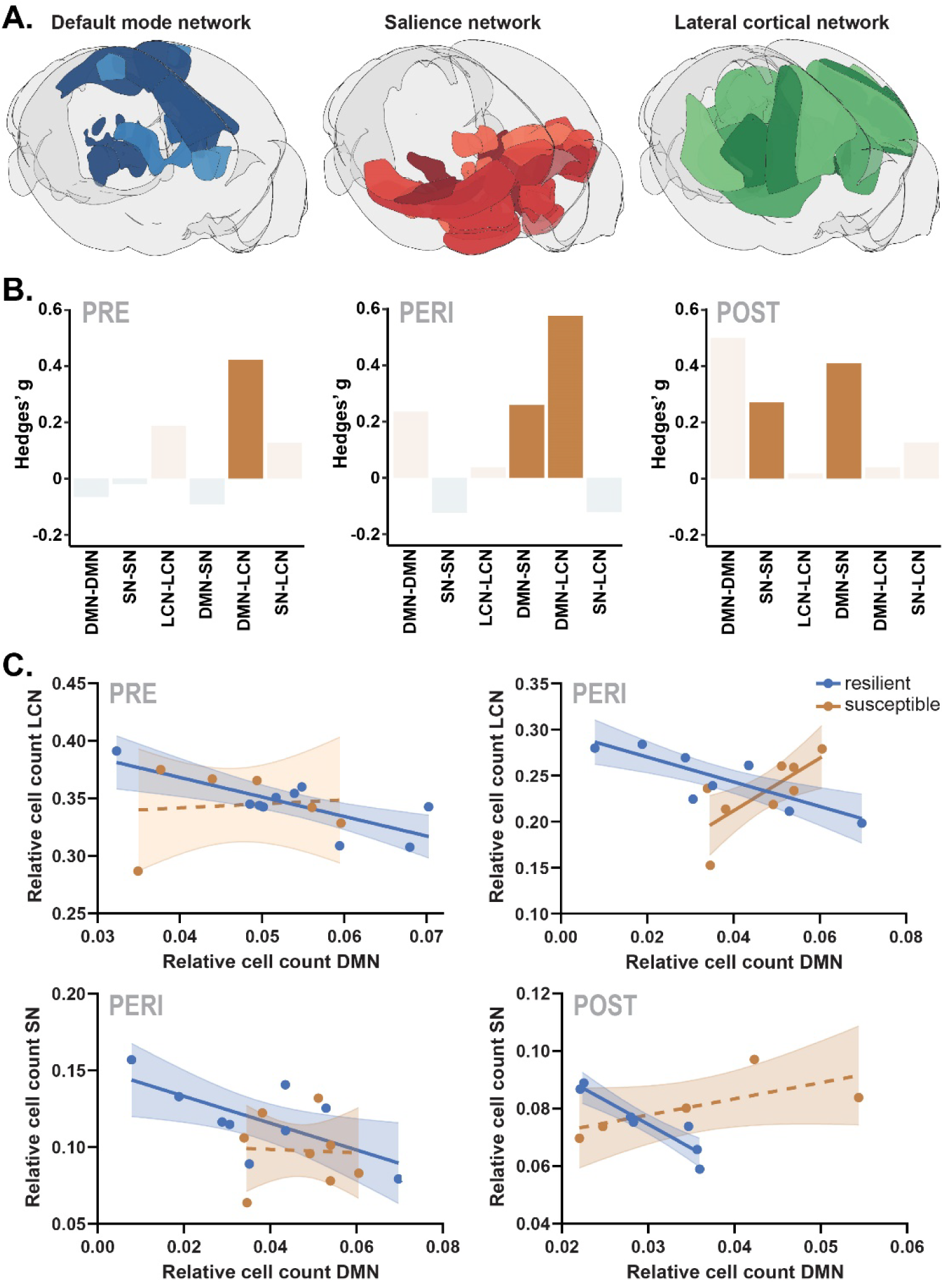
Differences in correlations of neuronal activity, indicative of inter-regional functional connectivity, within and across large-scale neural networks of susceptible vs resilient animals. Plots are based on Pearson correlation coefficients between regions in the default mode network (DMN, 7 regions), salience network (SN, 13 regions), and lateral cortical network (LCN, 11 regions, see Table S6) (**A**). Hedges’ g for the difference susceptible > resilient in network correlations of mice pre-, peri-, and post-stress are shown (**B**). To examine the observed differences in cross-network correlations between resilient and susceptible groups, total activity of each of the networks (i.e., the normalized total cell count across all the network’s subregions) was extracted and correlated among networks (**C**). Opaque bars in panel B indicate the differences in correlations where the 90% confidence interval does not cross or contain 0. Solid lines in panel C indicate correlations of which the 90% confidence interval does not cross or contain 0.

Except for previously reported elevations and reductions in isolated regions within these networks (Fig. 2), overall neuronal activity did not differ across susceptible and resilient mice in these networks in any of the cohorts (Fig. S3). We continued investigating potential differences between susceptible and resilient mice in terms of the functional connectivity between and within the networks by comparing cross-subject correlations in the activity of subregions part of these networks. To obtain estimates of overall intra- and inter-network connectivity, all correlations of individual subregions within each network (Fig. S4) were averaged to obtain a single correlation value. These analyses revealed that pre-stress, susceptible mice displayed increased correlations between the DMN and LCN (g_sus>res_ = 0.42 [0.16, 0.69], Fig. 4b) compared to resilient mice. A similar increase in DMN-LCN correlations was observed in susceptible mice at the peri-stress timepoint (g_sus>res_ = 0.58 [0.31, 0.84]). To investigate these differences further, total normalized cell counts within each of the networks were extracted and correlated with each other (Fig. 4c). DMN-LCN correlations observed both pre- (linear regression coefficient (lrc)_sus>res_ = 0.92 [0.16, 1.68]) and peri-stress (lrc_sus>res_ = 1.53 [0.91, 2.14]) differed between groups. Intriguingly, this difference seemed caused by the expected negative correlations in the resilient animals (pre: r = -0.80 [-0.96, -0.28], peri: -0.80 [-0.95, -0.41]), that were absent (pre: r = 0.13 [-0.77, 0.86]) or even positive (peri: r = 0.74 [0.21, 0.93]) in susceptible mice (Fig. 4c). Peri-stress, susceptible animals furthermore showed stronger correlations between the DMN and SN than their resilient counterparts (g_sus>res_ = 0.26 [0.01, 0.50]). This finding was also observed in the post-stress cohort (g_sus>res_ = 0.41 [0.16, 0.65]). Analyses of the extracted total normalized cell counts within the DMN and SN (Fig. 4c) confirmed that DMN-SN correlations differed between groups both peri-stress (lrc_sus>res_ = 1.00 [0.18, 1.83]) and post-stress (lrc_sus>res_ = 1.53 [0.68, 2.39]). These differences were related to the expected overall negatively correlated DMN-SN activity within the resilient groups (peri: r = -0.66 [-0.90, -0.13], post: r = -0.93 [-0.99, -0.68]), which was not observed in susceptible groups (peri: r = -0.05 [-0.66, 0.60], post: r = 0.70 [-0.28, 0.97]). Post-stress, stronger correlations were also observed between regions within the SN (g_sus>res_ = 0.27 [0.01, 0.53]) in susceptible vs resilient animals, indicative of greater intra-network connectivity. As such, our findings implicate deviant DMN-LCN connectivity as risk factor for the development of behavioral symptoms following stress exposure, whereas susceptible mice seem to acquire stronger DMN-SN and intra-SN connectivity over the course of stress exposure and recovery, respectively.

## Discussion

To better understand the neurobiological basis of stress-related psychopathology, it is essential to differentiate between adaptive and maladaptive stress responses^4,6,7^. In this study, we employed a mouse model to contrast mice that were either resilient or susceptible to the behavioral consequences of stress exposure. Rather than focusing on individual behaviors, we used a composite behavioral outcome score that incorporated multiple PTSD-like symptoms to better model the clinical situation^15^, where PTSD diagnosis involves a range of symptoms leading to a highly heterogeneous patient population^27^. We also observed significant variability in individual behavioral measures, both within and across experimental cohorts. Despite this behavioral heterogeneity, the emergence of consistent group differences confirmed the presence of shared neural correlates, which were replicated across cohorts. Some differences seemed to reflect pre-existing risk factors, such as hyperactivity in the lateral orbital area, while others, like hyperactivity in the dorsal retrosplenial area, emerged during stress exposure and were sustained post-stress. The most pronounced differences between susceptible and resilient mice were observed in the peri-stress period, where susceptible mice showed increased activity in sensory and memory-related regions, including the retrosplenial areas and subiculum, and reduced activity in the olfactory areas, ventral agranular insula, and dorsal striatum. Comparison of stress responding in susceptible vs resilient mice, defined by the alterations in neuronal activity observed peri-vs pre-stress, indicated that resilience was not simply linked to reduced stress responses but rather to a distinct pattern of neuronal activity. Resilient mice exhibited significant reductions in retrosplenial activity and increases in the agranular insula and ventral striatum. In contrast, susceptible mice showed heightened stress-induced activity in visual and memory-related areas, such as the temporal association areas, dentate gyrus, and pre- and postsubiculum. Additionally, susceptible mice displayed altered functional connectivity, with increased correlations between the default mode network (DMN) and lateral cortical network (LCN) pre- and peri-stress, elevated DMN-salience network (SN) connectivity peri- and post-stress and heightened intra-SN connectivity post-stress. A full discussion on all neural findings goes beyond the scope of this article, but we highlight the most relevant findings.

Susceptible mice displayed increased activity in the lateral orbital area under basal conditions both before and after stress exposure. Structural and functional alterations in the lateral orbitofrontal cortex have been reported before in PTSD^28^, most notably in relation to altered mood^29^ and emotional/threat responding^30,31^. Animal work has shown that increased lateral orbital activity relates to heightened passive stress coping behavior^32^, as well as impaired fear extinction, exaggerated renewal, and reduced contextualization of fear^33^. Our findings propose that resting activation in the lateral orbital area may be a predisposing risk factor for stress susceptibility. Research in healthy volunteers linking lower amplitude of low-frequency fluctuations in resting activity in the left orbitofrontal cortex to higher psychological resilience^34^ is in line with this. Notably, early-life stress has been associated with reduced orbitofrontal volumes^35^, which could potentially contribute to the increased risk it posits on the development of stress-related psychopathology later in life^36^.

Heightened activity in the retrosplenial areas emerged during the stress period in susceptible mice and persisted post-stress. The retrosplenial cortex, known for its role in contextual memory, serves as a hub region that integrates and coordinates information flow, facilitating both the acquisition and time-independent retrieval of contextual memories^37^. Supporting this, stimulation of the neural ensembles activated in the retrosplenial cortex during contextual learning has been found sufficient to induce contextual fear memory retrieval^38^. Moreover, the retroplenial cortex has been linked to self-referential processing and the retrieval of episodic memories^39^. Increased activity of the retrosplenial cortex has been observed in PTSD patients upon re-exposure to trauma-related stimuli^40,41^, and has been proposed to play an important role in the intrusive reexperiencing of their trauma because of its role in associative learning and priming^40^. Our findings suggest that heightened retrosplenial activity first emerges around the time of stress or trauma exposure, proposedly altering trauma memory processing, which is supported by the association between retrosplenial activity shortly after exposure to traumatic film clips and later intrusion symptoms in healthy volunteers^42,43^. This proposes the retrosplenial cortex as a novel target for early intervention studies in PTSD.

In addition to basal pre- and post-stress assessments, we here performed unprecedented measurements of neuronal activity *during* the exposure to a severe stressor, a timepoint typically inaccessible in humans. By comparing pre- and peri-stress neuronal activity maps in resilient and susceptible groups, we could reconstruct the neural correlates of adaptive vs maladaptive stress responding, respectively. Overall, both groups showed increased activation in visual areas, agranular insula, piriform cortex, ventral dentate gyrus, entorhinal areas, subiculum, and medulla during the stress period, while activity decreased in the primary motor area, primary somatosensory areas, dorsal and ventral thalamic regions, and midbrain areas. However, resilient mice exhibited marked reductions in retrosplenial activity and increases in the agranular insula and ventral striatum, suggesting an adaptive, active response to stress. While the exact implications of these changes remain speculative, previous research has shown that activation of dopaminergic projection neurons to the nucleus accumbens during stress promotes resilience and active coping behaviors in social defeat^44^, which seems in line with the increased ventral striatal activity. In contrast, susceptible mice displayed most dominant stress-related activity increases in visual areas as well as memory-related regions; the temporal association areas, dentate gyrus and pre- and postsubiculum. Excessive activity in visual processing regions has been reported before for PTSD^45^, and has been proposed to confer risk for developing stress-related psychopathology by enhancing the reactivation of traumatic memories in response to sensory cues^46,47^. Supporting this, activity in visual areas (and medial temporal lobe regions) during the encoding of trauma films has been linked to flashbacks in healthy volunteers^48^, and visual cortex inhibitory stimulation was found to reduce the emotional intensity of later intrusions^49^. Peri-trauma (i.e., 2 weeks post-trauma) morphology of the visual cortex correlates with PTSD symptom severity^50^ and peri-trauma (2-4 months post-trauma) activation of the occipital lobe was associated with peritraumatic dissociation, a strong predictor of PTSD^51^, further supporting the link between aberrant visual processing during stress/trauma exposure and risk for disease. Our data are the first to show such association in an animal model for stress susceptibility, and could be used to dissect the mechanistic underpinnings of this sensory sensitivity, being either the consequence of an exaggerated stress response, known to potentiate the perception of visual information^52,53^ or cause of it. The observed hyperactivity in memory-related regions, amongst which the subiculum, adds to the evidence of deviant memory processing of the stressful event^17^ in susceptible mice, and invites studies into the behavioral manifestations of these deviations.

Besides assessing differences in regional brain activity, we explored the functional connectivity within and across the DMN, SN, and LCN in the resilient and susceptible rodent brains based on the triple network model of psychopathology^20^. This model posits that deviations in dynamic cross-network interactions encode a wide range of pathologies. Specifically, in patients suffering from PTSD, a hyperactive and strongly intra-connected SN^54,55^ has been observed, as well as increased inter-network connectivity between the SN and DMN networks^55^. Both aberrations are thought to contribute to exaggerated threat-oriented and emotional self-reflective processing in PTSD. Here, we replicated these findings of enhanced intra-SN and DMN-SN connectivity post-stress exposure in our rodent model for stress susceptibility. Moreover, our observation of increased DMN-SN correlations peri-stress proposes this connectivity as early biomarker for stress susceptibility. Additionally, we found heightened correlations between DMN and LCN regions in susceptible animals compared to resilient ones, both before and during stress exposure. The DMN and ECN/LCN are typically anti-correlated^22,26,56,57^, but our findings indicated that only resilient mice displayed such negative correlation. It has been proposed that disrupted DMN-ECN coupling is associated with episodic memory deficits and could form the basis for intrusive trauma memory recollection^58^. While we did not find evidence of altered DMN-LCN coupling post-stress, our study does implicate peri-stress abnormalities in this circuitry in stress susceptibility, potentially by modulating memory processing of the stressful event.

Some limitations of the current work should be acknowledged. First, the approach of aligning downsampled images to a template brain is not ideally suited for analyzing very small regions, such as the basolateral amygdala or nucleus reuniens. As such, this study does not replace dedicated research focused on specific regions of interest but rather complements them. Additionally, we inferred network connectivity by calculating cross-subject activity correlations^59-61^ rather than examining changes in signal strength over time within individual animals, as would be possible by longitudinal measurements in fMRI studies.

Despite these limitations, this study demonstrates the power of whole-brain fluorescent labeling and 3D clearing techniques for investigating brain activity at precise timepoints during stress. Our approach provides an unprecedented view on pre-, peri- and post-stress neuronal activity, replicating key findings from human PTSD studies and uncovering new brain targets in stress susceptibility.

## Methods

### Animals

This study builds on a previous study that assessed amygdalar neuronal activity in animals susceptible to PTSD-like symptomatology^62^. While the current study has a broader aim and employs different techniques, the biological samples that have been analyzed were obtained from the same animals as were used in that study targeting the detailed assessment of neuronal activity in amygdalar subregions. The animal cohorts were part of three separate experiments: cohort 1 (n = 48) to assess brain-wide neuronal activity under resting (home cage) conditions pre-stress, cohort 2 (n = 44) to assess neuronal activity during (peri) stress exposure, and cohort 3 (n = 48) to assess neuronal activity under resting (home cage) conditions post-stress. Heterozygote ArcCreER^T2^xROSA offspring, referred to as ArcTRAP mice, were generated from crossing two founder mouse lines: ArcCreER^T2^ (B6.129(Cg)-*Arc^tm1.1(cre/ERT2)Luo^*/J, Jax strain ID 021881) and conditional tdTomato (B6.Cg-*Gt(ROSA)26Sor^tm9(CAG-tdTomato)Hze^*/J, Ai14, Jax strain ID 007909). These were purchased from The Jackson Laboratory and bred as described before^9^. The ArcTRAP genetic construct allows *Arc*-expressing (i.e., active) neurons to be labeled by the fluorescent protein tdTomato in a 48-hour time window after injection with the compound tamoxifen. Only male mice were used for this study, as this PTSD model^10,63^ has only been validated in males. Mice were group housed (3-4 mice per cage) in individually ventilated cages on a reverse 12 h light/dark cycle (09:00 - 21:00 h) at the Central Animal Facility of the Radboud University Nijmegen, The Netherlands, according to institutional guidelines. Food and water were provided *ad libitum*. Unless otherwise stated, behavioral testing was performed during the animal’s active phase (i.e., the dark) between 13.00 - 18.00 h. The experimental protocols were in line with international guidelines, the Care and Use of Mammals in Neuroscience and Behavioral Research (National Research Council 2003), the principles of laboratory animal care, as well as the Dutch law concerning animal welfare and approved by the Central Committee for Animal Experiments, Den Haag, The Netherlands (project ID: 2015-0090).

### General procedure

All mice were exposed to a stress induction paradigm (Fig. 1a) as described before^10,62,63^. Mice were exposed to a severe stressor (intense, unpredictable foot shocks) followed by a less severe trigger event (mild, predictable foot shocks) the next day. This trigger event is necessary for causing the long-term behavioral phenotype that is observed in susceptible mice in this model^10^, and can be compared to the human situation in which exposure to stressors and previous traumas predispose individuals to showing abnormal stress responses to later experiences^64^. After the repeated stress exposure and a week of recovery, mice were subjected to a set of behavioral tests over the course of two weeks to assess stress susceptibility by the assessment of PTSD-like symptoms. One week after the final behavioral test, mice were re-exposed to a fear-related context for 10 minutes and sacrificed by perfusion-fixation 90 minutes later (data not included here).

### Tamoxifen

All mice were injected with tamoxifen to induce fluorescent labeling of all *Arc*-expressing neurons at different time points during the protocol. Mice in the pre-stress cohort were injected with tamoxifen on day -3 – four days before the stressor – to label active neurons under basal home cage conditions. Mice in the peri-stress cohort were injected on the morning of day 1 – seven hours before the stressor – to label peri-stress activated neurons. Mice in the post-stress cohort were injected on day 19 – eighteen days after the initial stressor and four days before sacrifice – to label basal post-stress active neurons under home cage conditions. Tamoxifen was dissolved in a 10% ethanol / corn oil solution at a concentration of 10 mg/mL by overnight sonication and stored at -20°C until further use. Solutions were heated to body temperature and intraperitoneally injected at a dosage of 150 mg/kg to induce activity-dependent neuronal labeling.

### Stress protocol

Mice were individually placed in *Context A* boxes, in which they received 14 1 second 1.0 mA shocks (the ‘stressor’) over 85 minutes in variable intervals. For this, mice were first moved to the dark experimental room in groups of two to three animals in dark carton boxes before being placed in the fear-conditioning boxes, which were connected to a shock generator (Campden Instruments). *Context A* consisted of a black, triangular shaped Plexiglas box with a steel grid and metal tray. The boxes were sprayed with 1% acetic acid, not illuminated and 70 dB background noise was presented.

On the second day, mice were individually placed in *Context B* boxes, in which they received 5 1 second shocks of 0.7 mA over a period of five minutes (the ‘trigger’), presented over fixed intervals. For this trigger session, mice were moved to the 70 lux illuminated experimental room in see-through cages in groups of two to three animals. The *Context B* boxes contained curved white walls and a steel grid with a white tray underneath. The boxes were furthermore cleaned with 70% ethanol and during the session the house lights in the boxes were turned on. No background noise was presented.

Mice were allowed to recover for a week, after which their behavioral response to the stress protocol was assessed by testing for PTSD-like behavioral symptoms: impaired risk assessment (in the dark-light transfer test), increased anxiety (by marble burying), hypervigilance (by acoustic startle), impaired sensorimotor gaiting (by pre-pulse inhibition (PPI)), and disturbed circadian rhythm (by locomotor activity during the light phase)^10^.

### Behavioral assessments

#### Dark-light transfer test

On day 8 of the protocol, mice were tested in the dark-light transfer test. The test was executed in a box that was divided into a dark compartment (DC, 29 × 14 cm) and brightly illuminated (ca. 1100 lux) compartment (LC, 29 × 29 cm), connected by a retractable door. The mice were individually placed in the DC, and the door was opened to initiate a 5-min test session. Movement of the mice was recorded and scored automatically with Ethovision XT (Noldus IT). An additional area of 6 × 3 cm surrounding the opening of the LC was programmed into the software tracking measurements. Time spent in the LC as well as time spent in this ‘risk assessment’ zone were measured. Percentage risk assessment was calculated as the amount of time spent in the risk assessment zone as a percentage of total time spent in the LC.

#### Marble burying

On day 10, mice were individually placed in a 10 lux illuminated black open box (30 × 28 cm), containing a 5 cm deep layer of corn cobs, on top of which 20 marbles were centrally arranged in a 4 × 5 grid formation. Each mouse was placed in the corner of the box to initiate the task. Mice were videotaped for 25 min. Videos were scored by assessing the number of buried marbles after 25 min.

#### Startle response and pre-pulse inhibition

On day 12, mice were moved to the experimental room in their home cage and individually placed in small, see-through Plexiglas constrainers mounted on a vibration-sensitive platform inside a ventilated cabinet that contained two high-frequency loudspeakers (SR-LAB, San Diego Instruments). Movements of the mice were measured with a sensor inside of the platform. The pre-pulse inhibition test (PPI) started with an acclimatization period of 5 min, in which a background noise of 70 dB was presented, which was maintained throughout the entire 30-min session. Thirty-two startle cues of 120 dB, 40 ms in duration and with a random varying ITI (12–30 s), were presented with another 36 startle cues preceded by a 20 ms pre-pulse of either 75 dB, 80 dB or 85 dB. Sessions were scored by assessing the latency to peak startle amplitude of the 12 middle startle trials, and the pre-pulse inhibition, i.e., the percentage of startle inhibition response to the different pre-pulse stimuli [1 − (mean pre-pulse startle response/mean startle response without pre-pulse) × 100].

#### Light phase locomotion

Immediately after the pre-pulse inhibition test, mice were individually housed in Phenotyper cages (45 × 45 cm, Noldus) for 72 h while their locomotion was being recorded by an infrared-based automated system (EthoVision XT, Noldus). The first 24 h were considered habituation time and data were discarded. Total locomotion during the subsequent two light phases (21:00–09:00 h) was assessed.

### Behavioral categorization

In order to categorize mice as either stress susceptible or resilient, one compound measure was generated based on the five behavioral outcome scores. Mouse behavior on each of the tests was sorted, and the 20% of mice that had the lowest values were attributed 3 points for percentage risk assessment, 3 points for latency to peak startle amplitude, and 2 points for percentage PPI. Similarly, the 20% of mice showing the highest values were attributed 1 point for light locomotor activity and marble burying^63^. Points for each test were determined by factor analysis in which tests were clustered in three separate groups: (1) latency to peak startle amplitude and percentage risk assessment, (2) percentage PPI, and (3) marble burying and total light activity^10^. Ties in the marble burying test were resolved by also assessing the number of marbles buried after 15 minutes. The points per animal were tallied to generate an overall stress symptom score. Mice that had a total of four or more points (necessitating extreme behavior in multiple tests) were coined susceptible. Only mice that had zero points (indicating no abnormal behavior within any of the tests) were coined resilient. Notably, this approach allows for differential symptom profiles across susceptible mice.

### Re-exposure and sacrifice

On the final day of the experiment, day 23, mice were re-exposed to a fear-related context for 10 minutes to induce fear memory recall (data not included in this manuscript). No shocks were administered during this context re-exposure session. Mice were sacrificed 90 min post re-exposure under anesthesia (5% isoflurane inhalation followed by intraperitoneal injection of 200 μL pentobarbital) by perfusion with phosphate buffered saline (PBS) followed by 4% paraformaldehyde solution (PFA). The brains were surgically removed and post-fixed for 24 hours in 4% PFA, after which they were transferred to 0.1 M PBS with 0.01% sodium azide and stored at 4°C.

### Whole-brain immunostaining and clearing

Left hemispheres of susceptible and resilient animals of each cohort were processed following the iDISCO+ protocol for adult brains^11^. While most steps were followed in accordance with the existing protocol, two notable exceptions were made. First of all, no heparin was added to the PTwH buffer, as initial testing did not show any qualitative differences in clearing or staining upon addition or omission of this chemical. Furthermore, the samples were not incubated in 66% DCM / 33% methanol after the first dehydration series, as this was deemed unnecessary for proper delipidation of the brain, shortening the protocol by one day. In summary, the hemispheres were dehydrated using a methanol gradient, bleached in 5% H_2_O_2_ in methanol at 4°C overnight and subsequently rehydrated. As endogenous fluorescence was bleached during these steps, the hemispheres had to be immunolabelled for tdTomato. Additionally, cFos-positive cells were labelled to assess neuronal activity during stress-related context re-exposure (data not reported here). To do so, the hemispheres were permeabilized for 5 days at RT in permeabilization solution (PBS with 0.16% w/v Triton-X 100, 2.3% w/v glycine, 20% v/v DMSO), blocked for 4 days at 37°C (in blocking solution, PBS with 0.15% w/v Triton-X 100, 6% v/v donkey serum and 10% v/v DMSO) and then incubated with primary antibodies (rabbit anti-RFP, 1:750, 600-401-379, Rockland; guinea pig anti-cFos, 1:2.500, 226004, Synaptic Systems; in antibody buffer: PBS with 0.2% v/v Tween-20, 5% v/v DMSO and 3% v/v Donkey Serum) for 6 days at 37°C. Subsequently, brains were washed 5 × 1 h + 1 × overnight at RT, and incubated for 7 days with secondary antibodies (goat anti-rabbit Alexa-555, 1:200, A27039, Thermo Fisher; donkey anti-guinea pig, Alexa-647, 1:400, 706-605-148, Jackson ImmunoResearch; in antibody buffer) at 37 °C. Following 5 × 1 h + 1 × overnight washing at RT (PBS with 0.2% v/v Tween-20), samples were dehydrated in a methanol gradient and then incubated 2 x 1 h in 100% methanol, followed by 3 h in 66% DCM / 33% methanol and 2 × 15 min 100% DCM. Finally, the hemispheres were cleared in 100% dibenzyl ether (DBE, Sigma) in airtight glass vials. Brains were typically transparent within 2 h, and completely cleared overnight. Unfortunately, not all brain samples were cleared and/or stained successfully, meaning that some animals had to be excluded from further analysis. This exclusion encompassed 8 animals in the pre-stress cohort (2 resilient, 6 susceptible) and 8 animals in the post-stress cohort (5 resilient, 3 susceptible).

### Whole-brain imaging

The cleared hemispheres were imaged on a LaVision Ultramicroscope II light-sheet microscope, equipped with a NTK Photonics white-light laser and filter sets for 488 nm, 568 nm and 647 nm, imaged through a long-working distance objective (LaVision) at 1.1 × magnification (effective 2.2x, NA 0.1), and recorded with an Andor Neo 5.5 cooled sCMOS camera. Imaging was performed at 647 nm for capturing the cFos signal and at 555 nm to record the tdTomato signal. The emission light consisted of a triple light-sheet from the dorsal side of the brain at 0.54 NA, scanning at 2.95/2.95/3 µm x/y/z resolution (3 µm z-steps) with the “horizontal focus” method and 17-18 horizontal focus steps. The sample was imaged submerged in DBE in sagittal configuration, and the entire cerebrum fit inside a single field of view (x/y), with a typical brain producing ∼ 1600 z-planes of 3 µm each.

### Image preprocessing

The resulting image stacks were downsampled with a customized version of ClearMap 1^21^. Subsequently, each downsampled image stack was manually aligned with a template brain from the Allen Brain Atlas using the Bigwarp tool in FIJI, a landmark-based tool for deformable image alignment, by matching ca. 200 landmarks between each sample brain and the template brain. The experimenter was blinded to the experimental group. The tdTomato signal yielded sufficient spatial information for aligning the brains to the atlas, obviating the need for also capturing autofluorescence signal during imaging. See Fig. S5 for a high-resolution image of the tdTomato signal in one of the sagittal sections from a representative image stack. A warped version of the atlas was exported for each hemisphere, and overlain with the downsampled image stack of that particular brain for visual inspection of the quality of the alignment. Cell segmentation was performed in Arivis Vision4D software (Arivis GmbH, https://www.arivis.com) using the “Machine Learning Segmenter” plugin. For the purpose of this paper, only the tdTomato^+^ cells were considered. Cell coordinates and landmark coordinates were re-imported to ClearMap for mapping of the cells to the atlas and calculating cell counts per brain region. To guarantee that potential differences in cell counts would not be caused by inter-animal variation in signal quality, a brain mask was constructed that contained only areas of the brain covered by all image volumes and analysis was only performed therein (Fig. S6).

### Data analyses and statistics

#### Behavioral data

Behavioral data were analyzed using IBM SPSS Statistics 23. Data points deviating more than three inter-quartile ranges from the median were considered outliers and removed from further analysis. Hedges’ g was used as readout for differences in behavioral measurements (to be consistent with the approach implemented for the microscopy data). Effects were considered of relevance if the 90% confidence interval did not cross or contain 0.

#### Microscopy data

Preprocessing of the image stacks in ClearMap yielded two main outputs per animal: a heatmap of tdTomato^+^ cell density across the brain; and a dataset with raw tdTomato^+^ cell counts, allocated to 1241 brain regions as defined by the Allen Brain Atlas. Due to slight variations in staining, clearing and imaging quality, some variation existed in the total number of tdTomato^+^ cells detected. Importantly, total cell counts within each experimental cohort did not differ between susceptible and resilient mice (pre-stress: g_sus>res_ = 0.59 [-0.33, 1.44], peri-stress: g_sus>res_ = -0.25 [-0.93, 0.58], post-stress: g_sus>res_ = 0.27 [-0.73, 1.42], Fig. S7).

The heatmaps from all animals per group were resampled to 100 μm isotropic resolution, all signals were corrected for the total signal strength in the sample (i.e., total cell count, to account for differences in cell detection caused by variance in clearing quality), after which spatial smoothing was applied with a 200 μm full-width half-maximum kernel to enhance signal-to-noise ratio and to account for small mis-registrations. Heatmaps were loaded into R (‘oro.nifti’ package) and the effect size between susceptible and resilient animals was estimated using Hedges’ g (‘effectsize’ package) for every voxel. This is motivated by a) the mass univariate nature of the tests which are prone to false positives with p-value based inferences, and b) Hedges’ g being a better indicator over other standardized effect size indicators for group with n < 20.

Cell counts for each segregated region were corrected for the total cell count within the sample, and clustered into 92 larger anatomical regions (Table S4), to promote spatial accuracy and functional relevance. Then, Hedges’ g values were calculated per brain region to contrast susceptible and resilient animals per experimental cohort, or to contrast pre- and peri-stress labeling conditions per group. To model interactions, a multiple regression model was fitted to explain relative cell counts as a function of condition (pre- or peri-stress), group (resilient or susceptible), and their interactions. Normalized effects and their 90% confidence intervals for each factor were obtained using the parameters package for R. Effects were considered of relevance if the 90% confidence interval did not cross or contain 0.

Lastly, the DMN (7 regions), SN (13 regions), or LCN (11 regions) were defined (Table S5) based on previous viral tracer studies targeting these networks^24-26^, injecting virus in their core region (i.e., the anterior cingulate area, anterior insular area, and primary motor area, respectively) and assessing labels in projection regions. Brain regions identified as (i.e., labeled by) part of multiple brain networks (i.e., the claustrum, orbital area, prelimbic area, agranular insular area, frontal pole of the cerebral cortex, mediodorsal nucleus of the thalamus, substantia nigra compact part, central medial nucleus of the thalamus, caudoputamen, ventral medial nucleus of the thalamus, paracentral nucleus, secondary motor area, and gustatory areas) were assigned to the network to which they most strongly contributed (i.e., correlated with). Bivariate Pearson correlation coefficients were computed between the cell counts for these regions in susceptible and resilient animals separately and plotted as correlation heatmaps (Fig. S5). Hedges’ g values were estimated using the Pearson’s r values within or between networks as the statistical unit. To further explore group differences in network correlations, total cell counts per network were extracted and correlated between animals within groups.

### Data and code availability

The pre-processed data, consisting of cell counts per ROIs and 3-dimensional cell heat maps, as well as the code to reproduce the analyses, is available freely here: https://gitlab.socsci.ru.nl/preclinical-neuroimaging/stat_bart.

## Acknowledgements

This work was supported by a Veni (863.15.008) and Vidi (203.028) grant award to M.J.A.G.H. by the Netherlands Organization for Scientific Research.

## Author contributions

J.R.H., T.K., and M.J.A.G. planned and supervised the whole project, and together with B.C.J.D. designed the experiments. B.C.J.D., L.M., S.M., R.R. and A.B. conducted the behavioral testing and analyses, and B.C.J.D., M.N., H.M.M., R.M., J.G. and M.J.A.G.H. analyzed the iDISCO+ data. J.G. and M.J.A.G.H. supervised the iDISCO+ data analyses. B.C.J.D. and M.J.A.G. wrote the manuscript. All authors reviewed the manuscript.

