## Supplement for "The Neural Blueprint of Stress Susceptibility: Brain-wide neuronal activity associated with the consequences of stress"

**Supplementary Figures and Tables**

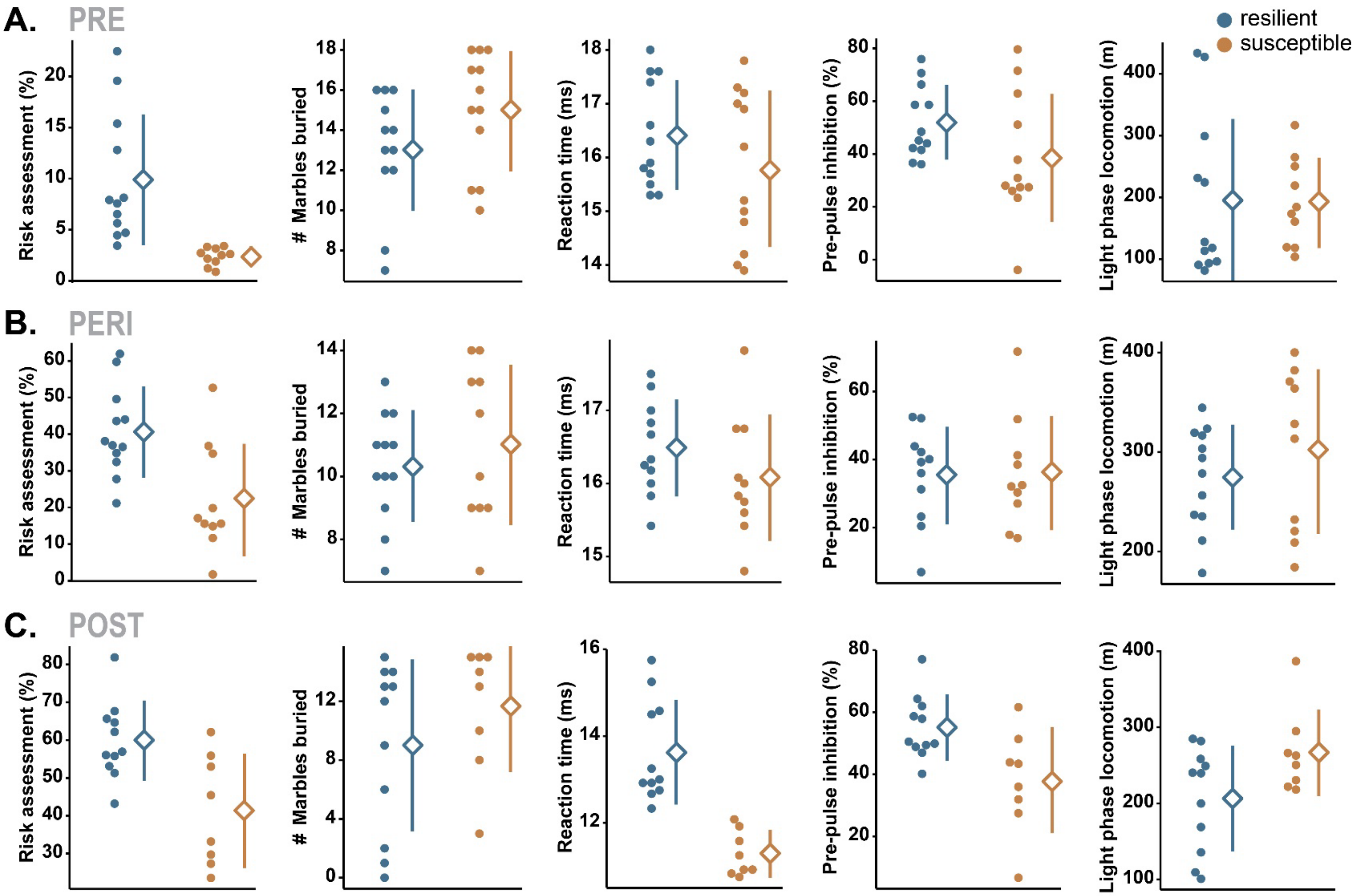

**Fig. S1.** Separate behavioral traits reflecting PTSD-like symptomatology across the three experimental cohorts in which neuronal activity was labeled pre-stress (**A**), peri-stress (**B**), and post-stress exposure (**C**). Individual data points are shown. Group means are depicted by a diamond, whereas the 90% confidence interval is indicated by the ends of the vertical error bars[1].

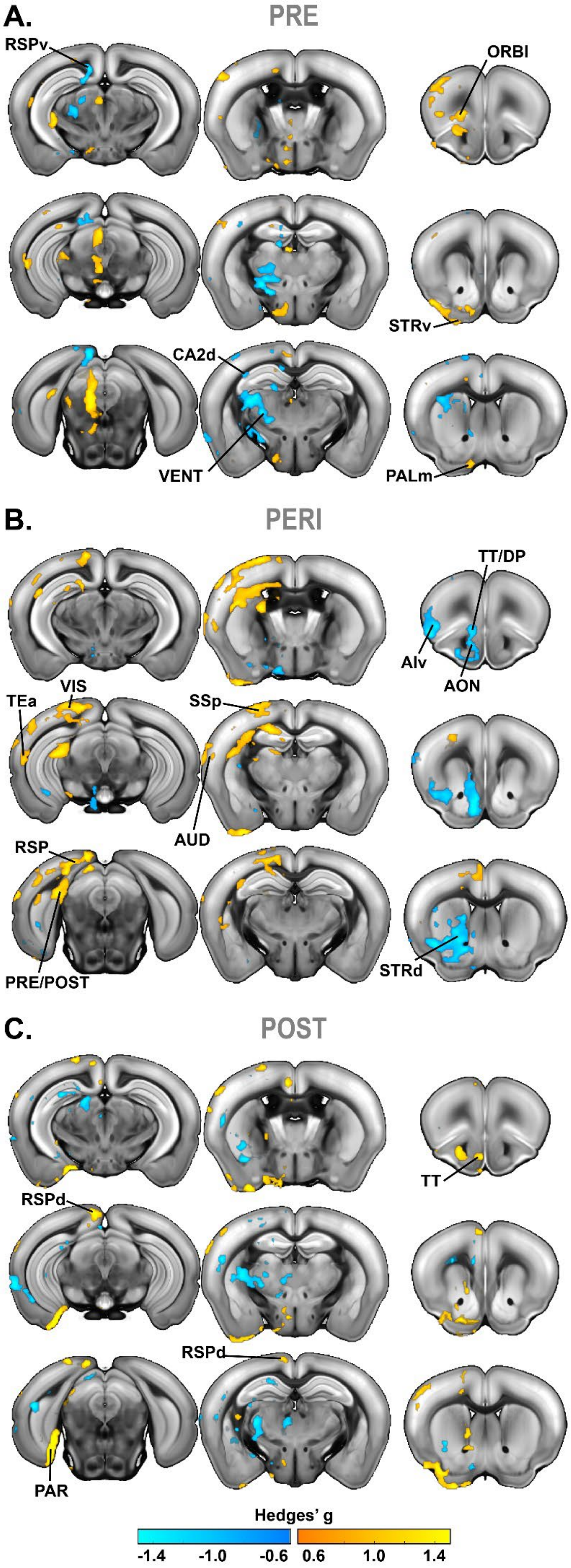

**Fig. S2.** Contrast maps generated by comparing neuronal activity (cell count) heat maps in susceptible vs resilient mice pre-stress (**A**), peri-stress (**B**) and post-stress exposure (**C**). Positive Hedges' g indicates higher density in susceptible mice. Maps are thresholded at 90% confidence intervals. AON: anterior olfactory nucleus, AIv: ventral agranular insular area, AUD: auditory areas, CA2d: dorsal hippocampal field CA2, DP: dorsal peduncular area, ORBl: lateral orbital area, PALm: medial pallidum, PAR: parasubiculum, POST: postsubiculum, PRE: presubiculum, RSPd: dorsal retrosplenial area, RSPv: ventral retrosplenial area, SSp: primary somatosensory area, STRd: dorsal striatum, STRv: ventral striatum, TEa: temporal association areas, TT: taenia tecta, VENT: ventral group of the dorsal thalamus, VIS: visual areas.

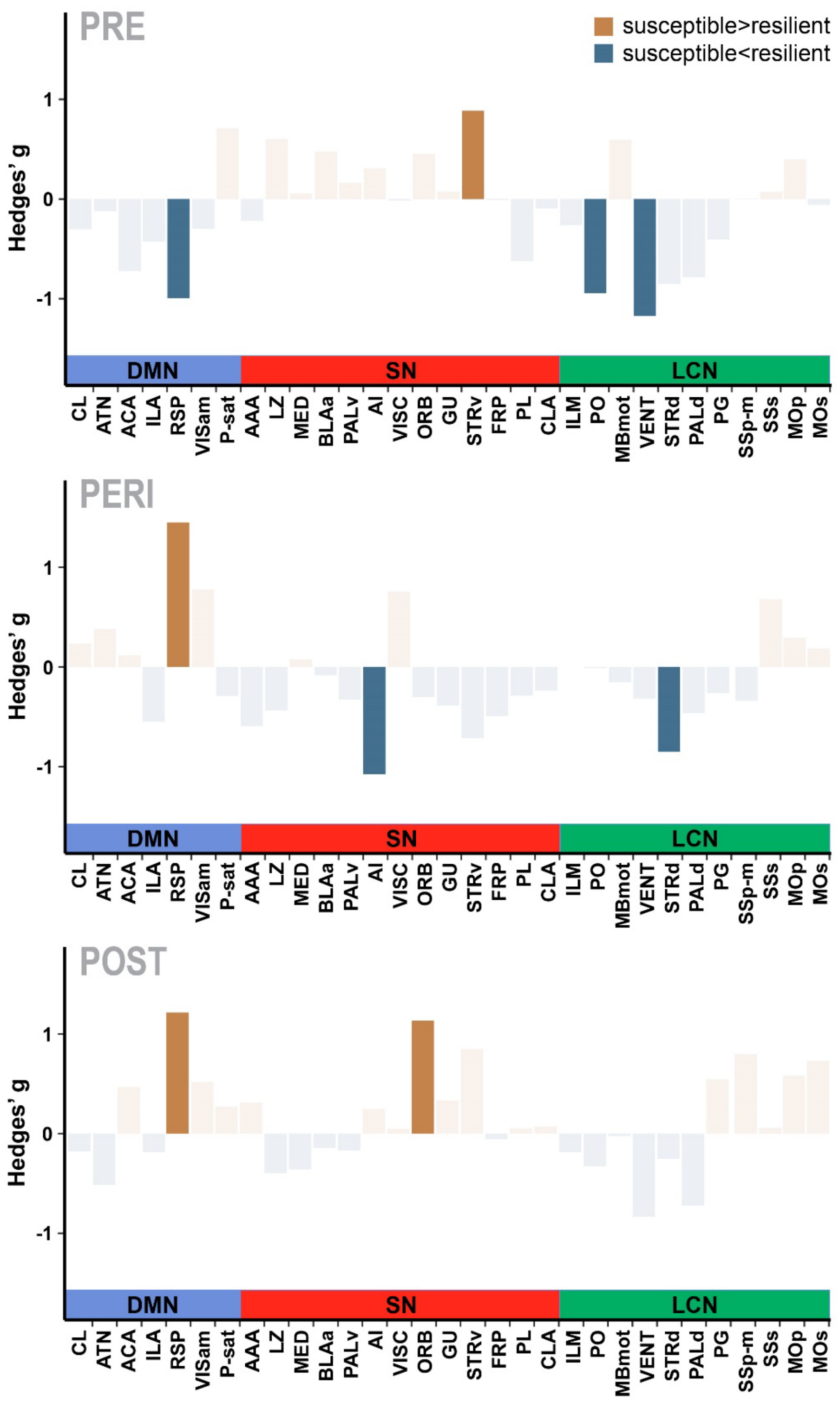

**Fig. S3.** Differences (susceptible > resilient) in relative neuronal activity (normalized cell counts) in the default mode network (DMN), salience network (SN), and lateral cortical network (LCN) between susceptible and resilient mice across experimental cohorts. Bars depicted in opaque colors indicate differences where the 90% confidence interval of the Hedges' g does not contain or cross zero; indicative of an effect. AAA: anterior amygdalar area, ACA: anterior cingulate area, AI: agranular insular area, ATN:

anterior nuclei of the dorsal thalamus, BLAa: anterior basolateral amygdalar nucleus, CL: central lateral nucleus of the thalamus, CLA: claustrum, FRP: frontal pole, GU: gustatory areas, ILA: infralimbic area, ILM: intralaminar nuclei of the dorsal thalamus, LZ: hypothalamic lateral zone, MBmot: midbrain, motor related, MED: medial group of the dorsal thalamus, MOp: primary motor area, MOs: secondary motor area, ORB: orbital area, PALd: dorsal pallidum, PALv: ventral pallidum, PG: pontine gray, PL: prelimbic area, PO: posterior complex of the thalamus, P-sat: pons, behavioral state related, RSP: retrosplenial area, SSp-m: primary somatosensory area, mouth, SSs: supplemental somatosensory area, STRd: dorsal striatum, STRv: ventral striatum, VENT: ventral group of the dorsal thalamus, VISam: anteromedial visual area, VISC: visceral area.

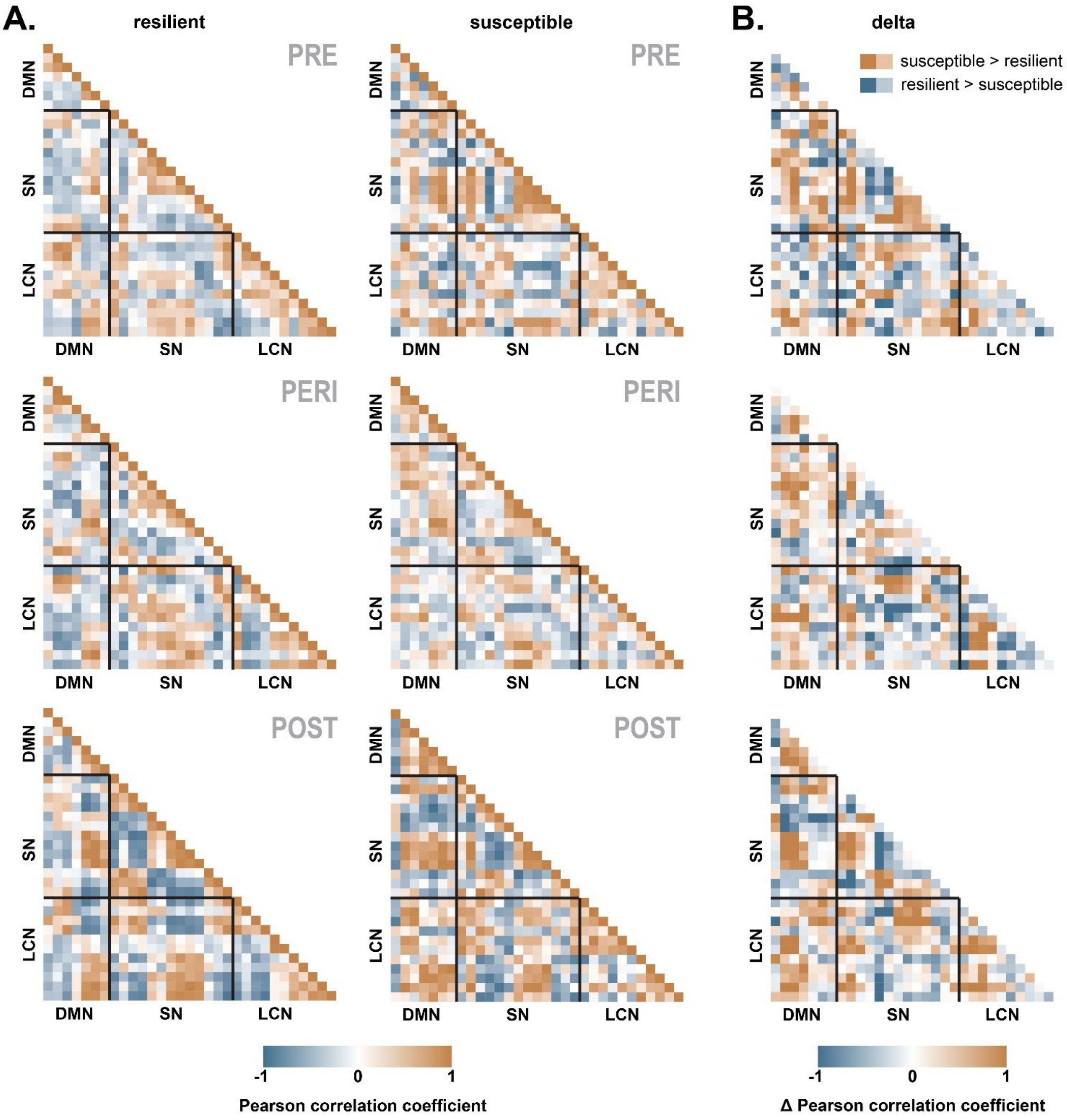

**Fig. S4.** Bivariate Pearson correlation heatmaps of the normalized cell counts of brain regions belonging to the default mode network (DMN), salience network (SN) and lateral cortical network (LCN) in susceptible and resilient animals (**A**), as well as their difference (susceptible > resilient) (**B**). The list of included brain regions can be found in Table S6.

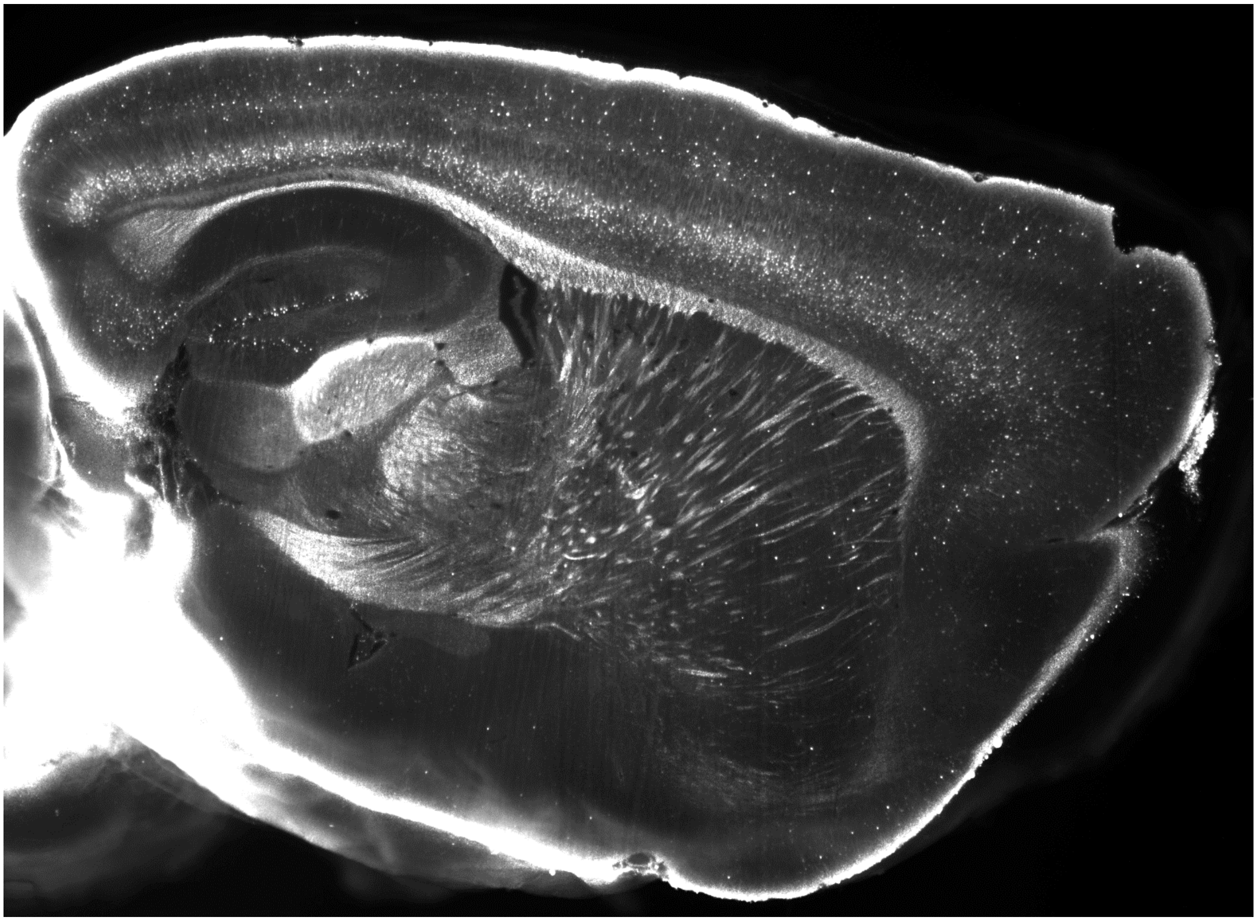

**Fig. S5.** High-resolution image of tdTomato$^+$ signal in a sagittal section from a representative 3D image stack.

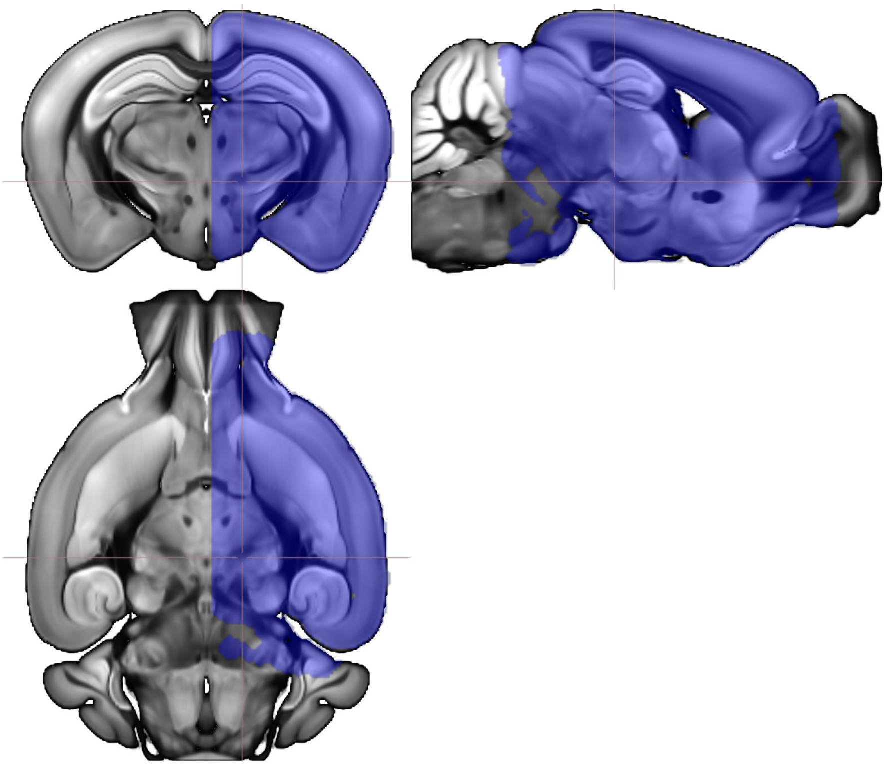

**Fig. S6.** Mask of included brain regions for the analyses of the iDISCO+ data.

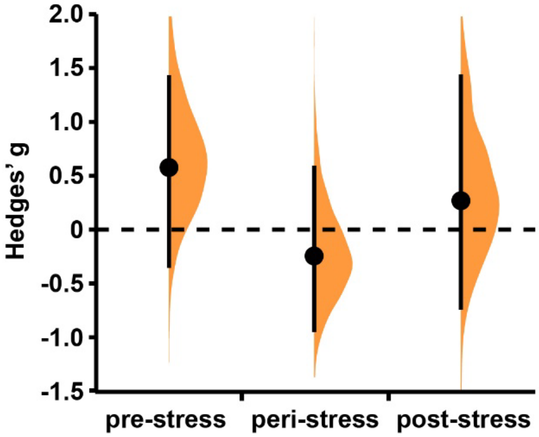

**Fig. S7.** Total cell counts detected following iDISCO+ in the three experimental cohorts did not differ between susceptible and resilient mice. Hedges' g for the difference susceptible > resilient is shown in Gardner-Altman estimation plots[1] for the total cell counts in the pre-, peri- and post-stress cohorts. Averaged total cell counts did differ across experimental cohorts. Whereas the total cell counts in the pre- and peri-stress cohorts did not differ from each other (pre (mean ± SD): 1.43 ± 0.28 x $10^6$ cells, peri: 1.80 ± 0.88 x $10^6$ cells, $g_{peri>pre}$ = 0.542 [-0.02, 0.96]), the total cell counts in the post-stress cohort (0.73 ± 0.28 x $10^6$ cells) were lower than in the other cohorts ($g_{pre>post}$ = 2.44 [1.63, 3.23] , $g_{peri>post}$ = 1.5 [0.96, 1.99]). Mean differences (susceptible – resilient) are depicted as a bootstrap sampling distribution; mean differences are indicated with a dot, whereas the 90% confidence intervals are indicated by the ends of the vertical error bars.

**Table S1. Differences between susceptible and resilient mice in behavioral traits reflecting PTSD-like symptomatology, as well as overall stress symptom score, over the three behavioral cohorts.**

| | Hedges' g [90% CI] |
|---|---|
| *Pre-stress* | |
| Risk assessment | -1.54 [-1.93, -1.12] |
| # Marbles buried | 0.66 [0.00, 1.32] |
| Reaction time to peak startle | -0.49 [-1.15, 0.20] |
| Pre-pulse inhibition | -0.68 [-1.36, 0.05] |
| Light phase locomotion | 0.33 [-0.40, 1.05] |
| Overall stress symptom score | 5.59 [4.32, 6.79] |
| *Peri-stress* | |
| Risk assessment | -1.33 [-2.16, -0.40] |
| # Marbles buried | 0.31 [-0.49, 1.01] |
| Reaction time to peak startle | -0.52 [ -1.29, 0.31] |
| Pre-pulse inhibition | 0.05 [-0.71, 0.78] |
| Light phase locomotion | 0.37 [-0.37, 1.13] |
| Overall stress symptom score | 5.07 [3.86, 6.80] |
| *Post-stress* | |
| Risk assessment | -1.45 [-2.32, -0.62] |
| # Marbles buried | 0.48 [-0.34, 1.13] |
| Reaction time to peak startle | -2.34 [-2.85, -1.75] |
| Pre-pulse inhibition | -1.25 [-1.90, -0.36] |
| Light phase locomotion | 0.92 [0.30, 1.52] |
| Overall stress symptom score | 8.82 [6.83, 11.10] |

Positive Hedges' g indicates susceptible > resilient.

**Table S2. List of clustered brain regions as analyzed and depicted in Fig. 2.**

| **Parent region** | **Subregion** | **Region #** | **Region** |
|---|---|---|---|
| Isocortex | | 1 | Frontal pole, cerebral cortex |
| | Motor areas | 2 | Primary motor area |
| | | 3 | Secondary motor area |
| | Somatosensory areas | 4 | Primary somatosensory area, nose |
| | | 5 | Primary somatosensory area, barrel field |
| | | 6 | Primary somatosensory area, lower limb |
| | | 7 | Primary somatosensory area, mouth |
| | | 8 | Primary somatosensory area, upper limb |
| | | 9 | Primary somatosensory area, trunk |
| | | 10 | Supplemental somatosensory area |
| | | 11 | Gustatory areas |
| | | 12 | Visceral area |
| | Auditory areas | 13 | Dorsal auditory area |
| | | 14 | Primary auditory area |
| | | 15 | Ventral auditory area |
| | Visual areas | 16 | Anterolateral visual area |
| | | 17 | Anteromedial visual area |
| | | 18 | Lateral visual area |
| | | 19 | Primary visual area |
| | | 20 | Posterolateral visual area |
| | | 21 | Posteromedial visual area |
| | Frontal areas | 22 | Anterior cingulate area, dorsal part |
| | | 23 | Anterior cingulate area, ventral part |
| | | 24 | Prelimbic area |
| | | 25 | Infralimbic area |
| | Orbital areas | 26 | Orbital area, lateral part |
| | | 27 | Orbital area, medial part |
| | | 28 | Orbital area, ventral part |
| | Agranular insula | 29 | Agranular insular area, dorsal part |
| | | 30 | Agranular insular area, posterior part |
| | | 31 | Agranular insular area, ventral part |
| | Retrosplenial areas | 32 | Retrosplenial area, lateral agranular part |

| | | | |
|---|---|---|---|
| | | 33 | Retrosplenial area, dorsal part |
| | | 34 | Retrosplenial area, ventral part |
| | | 35 | Temporal association areas |
| | | 36 | Perirhinal area |
| | | 37 | Ectorhinal area |
| Olfactory bulb | | 38 | Main olfactory bulb |
| | | 39 | Accessory olfactory bulb |
| | | 40 | Anterior olfactory nucleus |
| | | 41 | Taenia tecta |
| | | 42 | Dorsal peduncular area |
| | | 43 | Piriform area |
| | | 44 | Nucleus of the lateral olfactory tract |
| | | 45 | Cortical amygdalar area |
| | | 46 | Piriform-amygdalar area |
| | | 47 | Postpiriform transition area |
| Hippocampal formation | | 48 | Dorsal CA1 |
| | | 49 | Dorsal CA2 |
| | | 50 | Dorsal CA3 |
| | | 51 | Ventral CA1 |
| | | 52 | Ventral CA2 |
| | | 53 | Ventral CA3 |
| | | 54 | Dorsal DG |
| | | 55 | Ventral DG |
| | | 56 | Lateral entorhinal cortex |
| | | 57 | Medial entorhinal cortex |
| | | 58 | Ventral entorhinal cortex |
| | | 59 | Parasubiculum |
| | | 60 | Postsubiculum |
| | | 61 | Presubiculum |
| | | 62 | Subiculum |
| Cortical subplate | | 63 | Cortical subplate |
| | | 64 | Amygdala |
| Striatum | | 65 | Dorsal striatum |

| | | |
|---|---|---|
| | 66 | Ventral striatum |
| | 67 | Lateral septum complex |
| | 68 | Striatum-like amygdalar nuclei |
| Pallidum | 69 | Dorsal pallidum |
| | 70 | Ventral pallidum |
| | 71 | Medial pallidum |
| | 72 | Caudal pallidum |
| Thalamus | 73 | Ventral group of the dorsal thalamus |
| | 74 | Geniculate group of the dorsal thalamus |
| | 75 | Lateral thalamus |
| | 76 | Anterior thalamus |
| | 77 | Medial thalamus |
| | 78 | Interlaminar thalamus |
| | 79 | Geniculate thalamus |
| Hypothalamus | 80 | Periventricular zone |
| | 81 | Periventricular region |
| | 82 | Medial hypothalamus |
| | 83 | Lateral hypothalamus |
| Midbrain | 84 | Midbrain, sensory |
| | 85 | Midbrain, motor |
| | 86 | Periaqueductal gray |
| | 87 | Pretectal area |
| | 88 | Midbrain, behavioral state |
| Pons | 89 | Pons, sensory |
| | 90 | Pons, motor |
| | 91 | Pons, behavioral state |
| Medulla | 92 | Medulla |

**Table S3. Effect of stress exposure on relative neuronal activity as determined by the comparison of normalized cell counts in the pre- and peri-stress cohorts in resilient and susceptible mice.**

| | Hedges' g [90% CI] |
|---|---|
| *Resilient: Peri-stress > Pre-stress* | |
| Visual area, posterolateral | 1.33 [0.31, 2.30] |
| Agranular insular area, dorsal | 1.46 [0.42, 2.46] |
| ventral | 1.63 [0.56, 2.66] |
| posterior | 1.79 [0.69, 2.84] |
| Piriform area | 3.55 [2.02, 5.03] |
| Dentate gyrus, ventral | 1.09 [0.12, 2.04] |
| Entorhinal area, lateral | 2.14 [0.97, 3.28] |
| medial, dorsal zone | 1.70 [0.62, 2.74] |
| medial, ventral zone | 1.26 [0.26, 2.22] |
| Parasubiculum | 1.07 [0.10, 2.02] |
| Subiculum | 1.32 [0.31, 2.30] |
| Ventral striatum | 1.99 [0.85, 3.09] |
| Pons, sensory related | 1.76 [0.67, 2.82] |
| motor related | 1.03 [0.06, 1.97] |
| behavioral state related | 1.25 [0.25, 2.22] |
| Medulla | 1.84 [0.74, 2.91] |
| *Peri-stress < Pre-stress* | |
| Primary motor area | -1.67 [-2.71, -0.60] |
| Primary somatosensory area, barrel field | -1.30 [-2.28, -0.29] |
| mouth | -2.00 [-3.10, -0.86] |
| trunk | -1.03 [-1.97, -0.06] |
| Supplemental somatosensory area | -0.97 [-1.90, -0.01] |
| Dorsal auditory area | -1.00 [-1.93, -0.03] |
| Dorsal anterior cingulate area | -1.18 [-2.14, -0.19] |
| Retrosplenial area, dorsal | -1.61 [-2.63, -0.55] |
| ventral | -1.48 [-2.48, -0.44] |
| Dorsal thalamus, ventral group | -1.90 [-2.99, -0.78] |
| geniculate group | -2.21 [-3.35, -1.02] |
| lateral group | -3.15 [-4.52, -1.73] |

| | |
|---|---|
| anterior group | -1.57 [-2.59, -0.52] |
| medial group | -1.30 [-2.27, -0.29] |
| intralaminar nuclei | -1.26 [-2.22, -0.25] |
| Ventral thalamus, geniculate group | -3.64 [-5.15, -2.09] |
| Lateral hypothalamic area | -1.05 [-1.99, -0.08] |
| Pretectal region | -1.89 [-2.97, -0.77] |
| *Susceptible: Peri-stress > Pre-stress* | |
| Visceral area | 1.36 [0.21, 2.46] |
| Visual area, anterolateral | 1.65 [0.44, 2.46] |
| lateral | 2.59 [1.14, 3.99] |
| primary | 2.98 [1.42, 4.50] |
| posterolateral | 2.18 [0.84, 3.47] |
| Agranular insula, posterior | 1.96 [0.68, 3.19] |
| Retrosplenial area, lateral agranular | 1.58 [0.38, 2.73] |
| Temporal association areas | 1.29 [0.15, 2.38] |
| Ectorhinal area | 1.34 [0.19, 2.44] |
| Piriform area | 1.61 [0.41, 2.77] |
| Cortical amygdalar area | 1.65 [0.44, 2.82] |
| Dentate gyrus, ventral | 1.65 [0.44, 2.82] |
| Entorhinal area, lateral | 2.15 [0.82, 3.43] |
| medial, dorsal zone | 1.74 [0.51, 2.92] |
| medial, ventral zone | 1.44 [0.28, 2.56] |
| Postsubiculum | 2.10 [0.78, 3.36] |
| Presubiculum | 2.13 [0.81, 3.41] |
| Subiculum | 2.57 [1.12, 3.96] |
| Medulla | 1.27 [0.14, 2.36] |
| *Peri-stress < Pre-stress* | |
| Primary motor area | -1.98 [-3.22, -0.69] |
| Primary somatosensory area, mouth | -2.97 [-4.49, -1.41] |
| Infralimbic area | -1.63 [-2.78, -0.42] |
| Accessory olfactory bulb | -1.15 [-2.22, -0.04] |
| Dorsal striatum | -1.41 [-2.53, -0.25] |

| | |
|---|---|
| Dorsal thalamus, ventral group | -1.26 [-2.34, -0.13] |
| geniculate group | -2.59 [-3.98, -1.14] |
| lateral group | -1.65 [-2.82, -0.44] |
| intralaminar nuclei | -1.73 [-2.91, -0.50] |
| Ventral thalamus, geniculate group | -3.09 [-4.63, -1.49] |
| Lateral hypothalamic area | -2.05 [-3.30, -0.74] |
| Midbrain, motor related | -1.33 [-2.43, -0.19] |
| Pretectal region | -1.64 [-2.80, -0.43] |

**Table S4. Stress x group interaction effects in relative neuronal activity as determined by the comparison of normalized cell counts in susceptible and resilient mice in the pre- and peri-stress cohorts.**

| | Linear regression coefficient [90% CI] |
|---|---|
| *Susceptible (Peri-stress > Pre-stress) > Resilient (Peri-stress > Pre-stress)* | |
| Primary somatosensory area, lower limb | 1.20 [0.07, 2.33] |
| trunk | 1.46 [0.30, 2.62] |
| Dorsal auditory area | 1.19 [0.03, 2.35] |
| Visual area, anterolateral | 1.65 [0.59, 2.71] |
| lateral | 1.05 [0.13, 1.97] |
| primary | 1.28 [0.34, 2.22] |
| posterolateral | 1.25 [0.44, 2.06] |
| Retrosplenial area, lateral agranular | 2.02 [1.00, 3.04] |
| dorsal | 1.96 [0.99, 2.93] |
| ventral | 1.42 [0.41, 2.43] |
| Temporal association areas | 1.10 [0.01, 2.18] |
| Dentate gyrus, dorsal | 1.22 [0.05, 2.38] |
| ventral | 1.00 [0.03, 1.96] |
| Presubiculum | 1.56 [0.61, 2.50] |
| Postsubiculum | 1.46 [0.41, 2.51] |
| *Susceptible (Peri-stress > Pre-stress) < Resilient (Peri-stress > Pre-stress)* | |
| Agranular insula, dorsal | 1.32 [0.25, 2.39] |
| ventral | 1.12 [0.11, 2.13] |
| Anterior olfactory nucleus | 1.21 [0.05, 2.37] |
| Striatum, ventral | 1.21 [0.20, 2.22] |
| Pallidum, medial | 1.30 [0.12, 2.47] |
| Periventricular region | 1.30 [0.08, 2.51] |

**Table S5. Anatomical definitions of the default mode, salience and lateral cortical network.**

| Network | Region # | Region | Abbreviation |
|---|---|---|---|
| Default mode network | 1 | Central lateral nucleus of the thalamus | CL |
| | 2 | Anterior nuclei of the dorsal thalamus | ATN |
| | 3 | Anterior cingulate area | ACA |
| | 4 | Infralimbic area | ILA |
| | 5 | Retrosplenial area | RSP |
| | 6 | Anteromedial visual area | VISam |
| | 7 | Pons, behavioral state related | P-sat |
| Salience network | 8 | Anterior amygdalar area | AAA |
| | 9 | Hypothalamic lateral zone | LZ |
| | 10 | Medial group of the dorsal thalamus | MED |
| | 11 | Basolateral amygdalar nucleus, anterior part | BLAa |
| | 12 | Pallidum, ventral region | PALv |
| | 13 | Agranular insular area | AI |
| | 14 | Visceral area | VISC |
| | 15 | Orbital area | ORB |
| | 16 | Gustatory areas | GU |
| | 17 | Striatum, ventral region | STRv |
| | 18 | Frontal pole, cerebral cortex | FRP |
| | 19 | Prelimbic area | PL |
| | 20 | Claustrum | CLA |
| Lateral cortical network | 21 | Intralaminar nuclei of the dorsal thalamus | ILM |
| | 22 | Posterior complex of the thalamus | PO |
| | 23 | Midbrain, motor related | MBmot |
| | 24 | Ventral group of the dorsal thalamus | VENT |
| | 25 | Striatum, dorsal region | STRd |
| | 26 | Pallidum, dorsal region | PALd |
| | 27 | Pontine gray | PG |
| | 28 | Primary somatosensory area, mouth | SSp-m |
| | 29 | Supplemental somatosensory area | SSs |

| | | | |
|---|---|---|---|
| | 30 | Primary motor area | MOp |
| | 31 | Secondary motor area | MOs |
